# Evolution of Ty1 copy number control in yeast by horizontal transfer of a *gag* gene

**DOI:** 10.1101/741611

**Authors:** Wioletta Czaja, Douda Bensasson, Hyo Won Ahn, David J. Garfinkel, Casey M. Bergman

## Abstract

Insertion of mobile DNA sequences typically has deleterious effects on host fitness, and thus diverse mechanisms have evolved to control mobile element proliferation across the tree of life. Mobility of the Ty1 retrotransposon in *Saccharomyces* yeasts is regulated by a novel form of copy number control (CNC) mediated by a self-encoded restriction factor derived from the Ty1 *gag* capsid gene that inhibits virus-like particle function. Here, we survey a panel of wild and human-associated strains of *S. cerevisiae* and *S. paradoxus* to investigate how genomic Ty1 content influences variation in Ty1 mobility. We observe high levels of mobility for a canonical Ty1 tester element in permissive strains that either lack full-length Ty1 elements or only contain full-length copies of the Ty1’ subfamily that have a divergent *gag* sequence. In contrast, low levels of canonical Ty1 mobility are observed in restrictive strains carrying full-length Ty1 elements containing canonical *gag*. Phylogenomic analysis of full-length Ty1 elements revealed that Ty1’ is the ancestral subfamily present in wild strains of *S. cerevisiae*, and that canonical Ty1 in *S. cerevisiae* is a derived subfamily that acquired *gag* from *S. paradoxus* by horizontal transfer and recombination. Our results provide evidence that variation in the ability of *S. cerevisiae* and *S. paradoxus* strains to repress canonical Ty1 transposition *via* CNC is encoded by the genomic content of different Ty1 subfamilies, and that self-encoded forms of transposon control can spread across species boundaries by horizontal transfer.

## Introduction

Retrotransposons are mobile genetic elements that transpose *via* an RNA intermediate and impact genome size, structure, function and molecular evolution in diverse eukaryotic lineages (Chenais et al. 2012; Mita and Boeke 2016). The budding yeast *Saccharomyces cerevisiae* is a powerful model organism for studying retrovirus-like long terminal repeat (LTR) retrotransposons, with many fundamental aspects of retrotransposon biology initially characterized in this species (Voytas & Boeke, 2002; Curcio, Lutz & Lesage, 2015; Sandmeyer, Patterson & Bilanchone, 2015). The complete sequencing of the yeast genome provided the first insight into organization and evolution of retrotransposons at the genomic scale (Goffeau et al. 1996; Jordan and McDonald 1998; Kim et al. 1998; Jordan and McDonald 1999a; Promislow et al. 1999). More recently, advances in sequencing technologies and bioinformatics have provided unprecedented opportunities to investigate the evolutionary dynamics and consequences of transposition across yeast populations and species (Neuveglise et al. 2002; Liti et al. 2009; Carr et al. 2012; Bleykasten-Grosshans et al. 2013; Menconi et al. 2013; Istace et al. 2017; Nelson et al. 2017; Yue et al. 2017; Bergman 2018; Peter et al. 2018).

The current assembly of the *S. cerevisiae* S288c reference strain contains sequences from six families of LTR retrotransposons, four that are active (Ty1, Ty2, Ty3, and Ty4) and two that are inactive (Ty5 and Ty3_1p) (Kim et al. 1998; Carr et al. 2012). At least 50 complete and over 400 partial Ty elements comprise 3.3% of the S288c reference assembly (Kim et al. 1998; Carr et al. 2012). The abundance of complete or partial Ty elements and their solo LTR derivatives varies significantly between *S. cerevisiae* strains, where relatively high Ty content is observed in lab strains such as S288c relative to wild strains (Wilke et al. 1992; Liti et al. 2005; Liti et al. 2009; Carr et al. 2012; Bleykasten-Grosshans et al. 2013). In general, full-length Ty element insertions are found to be polymorphic across strains, while most solo LTR insertions and a few truncated “relict” elements are fixed or found at high allele frequency (Carr et al. 2012; Bleykasten-Grosshans et al. 2013). Additionally, some Ty elements in *S. cerevisiae* have distinct subfamilies (e.g. the Ty1’ and Ty1/2 subfamilies of Ty1) (Jordan and McDonald 1998; Kim et al. 1998; Jordan and McDonald 1999b) or show evidence of recent horizontal transmission from other species (e.g. Ty2 and Ty3_1p) (Liti et al. 2005; Carr et al. 2012).

Ty1 is the most abundant retrotransposon family in the *S. cerevisiae* reference strain S288c (>30 full-length copies), and is both actively transcribed and transpositionally competent (reviewed in Curcio, Lutz & Lesage (2015)). The structure and replication of Ty1 elements resembles that of retroviruses. Ty1 consists of two partially overlapping open reading frames – *gag* (*TYA*) and *pol* (*TYB*) – flanked by LTRs. mRNA from full-length Ty1 elements serves as a template for both reverse transcription and translation of the proteins necessary for retrotransposition: the Gag capsid protein, protease, integrase, and reverse transcriptase. Ty1 RNA is specifically packaged into virus-like particles (VLPs) and serves as the template for reverse transcription into linear cDNA, which subsequently is imported into the nucleus as a protein/DNA complex using a nuclear localization signal present on integrase. Ty1 preferentially integrates near genes transcribed by RNA Polymerase III through an association between integrase and Pol III-complexes (Bridier-Nahmias et al. 2015; Cheung et al. 2016).

Because of the deleterious effects of most transposition events, eukaryotic hosts have evolved effective mechanisms to restrict the mobility or expression of transposons including RNAi, DNA methylation, and APOBEC proteins (Friedli and Trono 2015; Goodier 2016). Importantly, none of those systems operate natively in *S. cerevisiae* or its sister species, *S. paradoxus* (Drinnenberg et al. 2009). Instead, Ty1 mobility in *S. cerevisiae* and *S. paradoxus* is limited by a novel retroelement-directed restriction mechanism termed Copy Number Control (CNC) (Wilke et al. 1992; Garfinkel et al. 2003; Saha et al. 2015; Garfinkel et al. 2016; Ahn et al. 2017). CNC is defined as a decrease in Ty1 mobility when additional copies of the Ty1 element are present in the genome. CNC is mediated by the Ty1 restriction protein p22, which is a truncated version of Gag encoded by internally-initiated Ty1 transcripts (Saha et al. 2015). p22 interferes with a central function of the Ty1 capsid during VLP assembly and maturation, and thus is a potent self-encoded *trans*-dominant negative inhibitor of Ty1 retrotransposition (Nishida et al. 2015; Saha et al. 2015; Tucker et al. 2015). Ty1 inhibition by p22 bears striking similarities to host-encoded restriction factors that inhibit retrovirus assembly or uncoating (Tucker and Garfinkel 2016).

To date, Ty1 CNC has been studied in a very limited number of genetic backgrounds: first in *S. paradoxus* strain 337 (Garfinkel et al. 2003; Moore et al. 2004; Garfinkel 2005; Saha et al. 2015) and more recently in *S. cerevisiae* strain DJ12 (Ahn et al. 2017). Thus, it remains an open question at what level CNC operates in diverse lineages of *S. cerevisiae* and *S. paradoxus* that vary in their endogenous genomic Ty1 content. Here we use well-developed methods for detecting frequency of Ty1 movement (Curcio and Garfinkel 1991; Atwood et al. 1998; Curcio et al. 2007) to measure mobility of a “canonical” *S. cerevisiae* Ty1 tester element in a diverse panel of *S. cerevisiae* and *S. paradoxus* strains. Our results reveal that canonical Ty1 mobility varies substantially among strains in both species. We show that “permissive” strains with high Ty1 mobility can be converted to “restrictive” strains with low Ty1 mobility by experimentally introducing multiple canonical Ty1 elements into permissive genomes, implying that variation in mobility is due to variation in the strength of CNC rather than other differences in genetic background. Additionally, we investigated the genomic basis of variation in Ty1 mobility using whole genome PacBio long-read assemblies that yield complete sequence information of transposable elements in their native chromosomal locations (Khatri et al. 2017; Yue et al. 2017; Naseeb et al. 2018). By comparing Ty1 copy number and sequence composition with mobility frequency, we infer that restrictive strains in both *S. cerevisiae* and *S. paradoxus* contain full-length Ty1 elements with a canonical form of *gag*. In contrast, permissive strains either lack full-length Ty1 elements or only contain full-length elements from the Ty1’ subfamily that have a divergent *gag* sequence. Surprisingly, the reconstructed evolutionary history of full-length Ty1 elements in *S. cerevisiae* and *S. paradoxus* shows that the Ty1’ subfamily is the ancestral subfamily in *S. cerevisiae* found in wild lineages, while the canonical Ty1 family used in most functional studies is a highly-derived element found in human-associated strains. Furthermore, we discovered that the *gag* region of the canonical *S. cerevisiae* Ty1 element was acquired by horizontal transfer from an Old-World lineage of *S. paradoxus* followed by recombination onto a pre-existing ancestral Ty1’-like element. Our results demonstrate that intraspecific variation in the ability to repress transposition of the canonical Ty1 subfamily in *S. cerevisiae* is a consequence of horizontal transfer of a CNC-competent *gag* gene from a closely-related yeast species.

## Results

### Ty1 restriction varies across diverse isolates of *S. cerevisiae* and *S. paradoxus*

Because Ty1 CNC is mediated by a self-encoded factor (p22) and dependent on Ty1 genomic copy number, we hypothesized that variation in endogenous Ty1 genomic content may influence the strength of Ty1 CNC across *Saccharomyces* strains. To address this possibility, we first performed Southern analysis on a set of genetically-tractable haploid derivatives of 25 *S. cerevisiae* strains and 27 *S. paradoxus* strains from the *Saccharomyces* Genome Resequencing Project (SGRP) to understand how these strains vary in their Ty1 content (Cubillos et al. 2009; Liti et al. 2009). The probe for hybridization experiments in both species is derived from the *gag* region of the “canonical” Ty1-H3 element (Sup Fig 1A), a full-length Ty1 element isolated in *S. cerevisiae* as a *His^+^* reversion mutant that has been used in many pioneering studies on Ty1 structure and function (Boeke et al. 1985; Boeke et al. 1988). This analysis revealed substantial diversity across both *S. cerevisiae* and *S. paradoxus* strains in the number of Ty1 elements that share strong sequence similarity to canonical Ty1-H3 *gag* (Sup Fig 1B and Sup Fig 1C), with most strains having fewer Ty1 elements than the *S. cerevisiae* reference strain S288c. These results also revealed several strains that contained no elements with similarity to canonical Ty1 *gag*, consistent with the existence of multiple “Ty1-less” strains that lack full-length elements in *S. cerevisiae* and *S. paradoxus* (Wilke et al. 1992; Moore et al. 2004; Istace et al. 2017; Yue et al. 2017).

Next, we selected a diverse panel of ten *S. cerevisiae* and *S. paradoxus* SGRP strains with distinct Ty1 hybridization patterns (Figure 1A) to test for variation in the frequency of mobility using a canonical Ty1-H3 tester element marked with a *his3-AI* indicator gene (Curcio and Garfinkel 1991). We performed Ty1 mobility assays in seven *S. cerevisiae* strains (S288c, Y12, DBVPG6044, UWOPS83-787.3, YPS606, UWOPS05-227.2, L-1374) by introducing a *URA3*-based centromere plasmid containing a competent Ty1*his3-AI* element into haploid *MATα his3-Δ200hisG* SGRP strains. The frequency of His^+^ colony formation in this assay detects Ty1 mobility events from either *de novo* retrotransposition events or insertion events from a minor pathway where Ty1 cDNA undergoes homologous recombination with genomic or plasmid-borne Ty1 sequences (Sharon et al. 1994). For the three *S. paradoxus* strains tested (CBS432, N-44, YPS138), deletion of *HIS3* could not be achieved efficiently and thus Ty1 mobility assays were performed by first replacing the *KanMX* gene inserted at the *URA3* locus with *NatMX* in haploid *MATα* SGRP strains, then introducing a reporter plasmid containing Ty1*neo-AI*. The appearance of G418-resistant colonies in these *S. paradoxus* strains is a readout for retromobility that can be monitored by qualitative or quantitative assays similar to Ty1*his3-AI* reporter system (Curcio and Garfinkel 1991; Atwood et al. 1998; Curcio et al. 2007). These experiments revealed >50-fold differences in the mobility of canonical Ty1 across strains within both *S. cerevisiae* and *S. paradoxus* (Table 1). In both species, we observed “restrictive” strains with very low levels of canonical Ty1 mobility (*S. cerevisiae*: S288c, Y12, and DBVPG6044; *S. paradoxus*: CBS432, N-44). Likewise, we observed “permissive” strains in both species with canonical Ty1 mobility frequencies which were more than an order of magnitude higher than restrictive strains (*S. cerevisiae*: UWOPS05-787.3, YPS606, UWOPS05-227.2, and L1374; *S. paradoxus*: YPS138).

**Figure 1.**
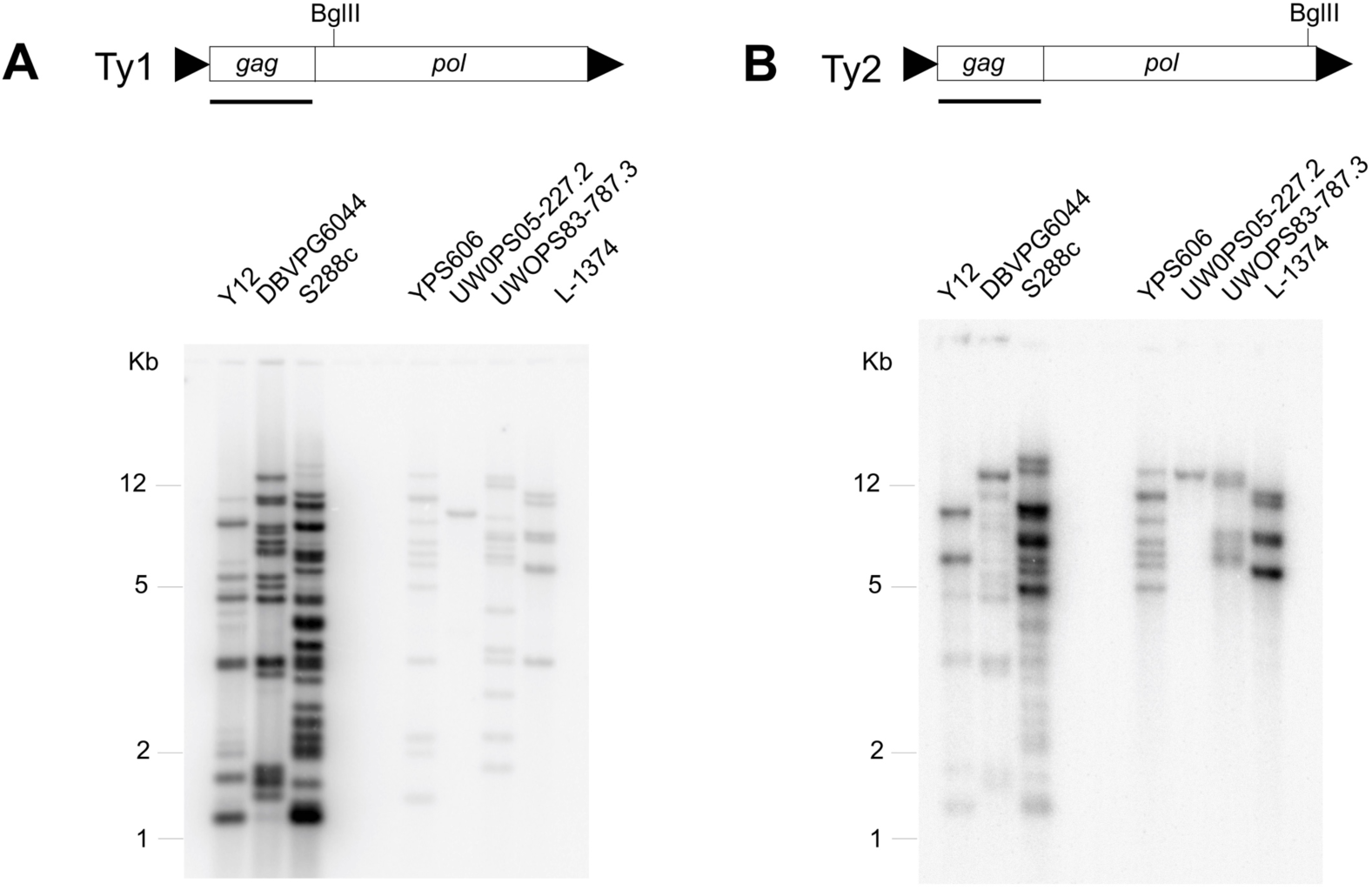
Southern blots of canonical Ty1-H3 and Ty2 *gag* hybridized to *S. cerevisiae* strains with mobility phenotypes. Southern blot results using radiolabeled (**A**) Ty1 and (**B**) Ty2 *gag* probes for *S. cerevisiae* strains with mobility data. Restrictive strains are shown on the left, and permissive strains are shown on the right of each Southern blot. Schematics of the Ty1-H3 and Ty2 elements, respectively, showing the *gag* and *pol* open reading frames (rectangles) and LTRs (arrowheads) are shown above each Southern blot. The probe used for Southern blots is obtained from the *gag* gene (underlined). The locations of the BglII restriction sites within Ty1-H3 and Ty2 *pol* regions are shown above the schematics. Identical membranes with the same DNA samples were hybridized in panels (A) and (B).

**Table 1.** Canonical Ty1-H3 mobility in a diverse panel of *S. cerevisiae* and *S. paradoxus* strains. Ty1 mobility frequency is defined as the number revertant colonies (His^+^ Ura^+^ colonies for Ty1*his3*-AI or G418^R^ colonies for Ty1*neo*-AI) colonies divided by the number of Ura^+^ colonies per ml of culture. Standard deviations were calculated from the number Ty1 mobility events detected per 1 ml culture. Because of differences in reporter constructs and selection systems, mobility frequencies can be compared across strains within species, but not across species. S.D. = standard deviation. N.D. = not determined.

| Strain | Species | Source | Origin | Ty1*his3*-AI<br>x10<sup>-6</sup><br>[S.D.] | Ty1*neo*-AI<br>x10<sup>-7</sup><br>[S.D.] | Ty1<br>mobility<br>phenotype |
| --- | --- | --- | --- | --- | --- | --- |
| S288c | *S. cerevisiae* | Lab | N. America | 1.6 [0.2] | N.D. | restrictive |
| DBVPG6044 | *S. cerevisiae* | Bili Wine | West Africa | 1.0 [0.17] | N.D. | restrictive |
| Y12 | *S. cerevisiae* | Sake | Japan | 5.6 [0.5] | N.D. | restrictive |
| UWOPS83-787.3 | *S. cerevisiae* | Wild | Bahamas | 78 [5.5] | N.D. | permissive |
| YPS606 | *S. cerevisiae* | Wild | N. America | 157 [13] | N.D. | permissive |
| UWOPS05-227.2 | *S. cerevisiae* | Wild | Malaysia | 458 [86] | N.D. | permissive |
| L-1374 | *S. cerevisiae* | Wine | Chile | 635 [79] | N.D. | permissive |
| CBS432 | *S. paradoxus* | Wild | Europe | N.D. | <0.6 | restrictive |
| N-44 | *S. paradoxus* | Wild | Far East Asia | N.D. | 0.9 [0.8] | restrictive |
| YPS138 | *S. paradoxus* | Wild | N. America | N.D. | 31 [7] | permissive |

Variation in Ty1 mobility across *Saccharomyces* strains could result from variation in the strength of CNC conferred by endogenous Ty1 elements or other differences in host genetic background. To determine whether variation in Ty1 mobility is more likely due to Ty1 CNC or a Ty1-independent host background effect, we over-expressed canonical Ty1-H3 from a plasmid to “populate” the genomes of three permissive *S. cerevisiae* strains (UWOPS05-227.2, L1374, and YPS606) with >8 canonical Ty1 elements, as estimated by Southern analysis (see Materials and Methods for details). As shown previously for *S. paradoxus* strain 337 (Garfinkel et al. 2003), we observed a >60-fold decrease in canonical Ty1 mobility in three *S. cerevisiae* strains populated with multiple canonical Ty1 elements when compared with their respective native parental strains (Table 2), with canonical Ty1 mobility in populated strains being on the same order as other native strains with restrictive phenotypes. We note that mobility data for native strains in Table 2 were from an independent set of experiments done in parallel with populated strains and thus differ slightly from the data in Table 1 for the same native strains. The ability for permissive strains to become restrictive with the addition of full-length copies of canonical Ty1 indicates that genetic background effects alone cannot explain variation in Ty1 mobility and is consistent with Ty1 CNC playing a major role in shaping variation in Ty1 mobility among yeast strains.

**Table 2.** Canonical Ty1-H3 mobility in natively permissive *S. cerevisiae* strains populated with canonical Ty1-H3 elements. Canonical Ty1 copy number in populated strains was estimated by Southern blot analysis as described previously (Garfinkel et al. 2003).

| Strain | Species | Ty1$his3-AI$ x10<sup>-6</sup> [SD] | Ty1 mobility phenotype |
| --- | --- | --- | --- |
| YPS606 | *S. cerevisiae* | 164 [8.5] | permissive |
| YPS606 + 8 canonical Ty1's | *S. cerevisiae* | 2.4 [0.12] | restrictive |
| UWOPS05-227.2 | *S. cerevisiae* | 529 [124] | permissive |
| UWOPS05-227.2 + 17 canonical Ty1's | *S. cerevisiae* | 1 [0.25] | restrictive |
| L-1374 | *S. cerevisiae* | 461 [163] | permissive |
| L-1374 + 20 canonical Ty1's | *S. cerevisiae* | 3 [0.35] | restrictive |

### The presence of full-length Ty1 elements is not sufficient to restrict Ty1-H3 mobility

To determine if variation in Ty1 mobility is influenced by the copy number or sequence of endogenous Ty1 elements, we generated ~100x whole-genome shotgun PacBio datasets and assembled genome sequences for the seven *S. cerevisiae* strains we assayed for Ty1 mobility. We integrated data from our *S. cerevisiae* PacBio assemblies with similar high quality PacBio genome assemblies from Yue *et al*. (2017) for the three strains of *S. paradoxus* with mobility data in our study (CBS432, N-44, YPS138). PacBio assemblies typically reconstructed complete chromosomes in single contigs (with the exception of chromosome XII which was broken at the highly repeated rDNA locus) and thus provide an essentially-complete catalogue of Ty content in yeast genomes. We identified Ty elements in these ten PacBio assemblies using a RepeatMasker-based strategy that classifies Ty elements as full-length, truncated, or solo LTR sequences based on the completeness of internal sequences in each predicted element (see Materials and Methods for details). Although our focus is on Ty1, we annotated all Ty families in these genomes to avoid potential misidentification, and because the similarity of solo LTRs from Ty1 and Ty2 does not allow their unambiguous assignment to either family (see also Yue *et al*. (2017)). Predicted numbers of full-length, truncated, or solo LTR sequences for Ty1 can be found in Table 3 and for all Ty families in Sup File 1. We focused on full-length elements in our analysis since they are most likely to have the complete set of functional sequences required for Ty1 gene expression and transposition.

**Table 3.** Ty1 content in a diverse panel of *S. cerevisiae* and *S. paradoxus* strains with Ty1 mobility phenotypes. Asterisks represent inclusion of two elements (*: Y12_f109; **: S288c_f486) that are recombinant between Ty1’ and canonical Ty1 *gag* and are classified according to the group from which the majority of their *gag* sequence is derived.

| Strain | Species | Ty1 mobility phenotype | Full-length Ty1 | Truncated Ty1 | Ty1/Ty2 solo LTRs | Full-length with canonical Ty1 *gag* | Full-length with Ty1' *gag* |
| --- | --- | --- | --- | --- | --- | --- | --- |
| S288c | *S. cerevisiae* | restrictive | 38 | 2 | 161 | 35 | 3** |
| DBVPG6044 | *S. cerevisiae* | restrictive | 19 | 5 | 269 | 19 | 0 |
| Y12 | *S. cerevisiae* | restrictive | 19 | 2 | 158 | 8* | 11 |
| UWOPS83-787.3 | *S. cerevisiae* | permissive | 7 | 3 | 158 | 0 | 7 |
| YPS606 | *S. cerevisiae* | permissive | 3 | 2 | 147 | 0 | 3 |
| UWOPS05-227.2 | *S. cerevisiae* | permissive | 0 | 2 | 185 | 0 | 0 |
| L-1374 | *S. cerevisiae* | permissive | 0 | 1 | 155 | 0 | 0 |
| CBS432 | *S. paradoxus* | restrictive | 8 | 4 | 264 | 8 | 0 |
| N-44 | *S. paradoxus* | restrictive | 2 | 3 | 235 | 2 | 0 |
| YPS138 | *S. paradoxus* | permissive | 0 | 1 | 232 | 0 | 0 |

The total number of full-length Ty1 elements varies substantially across the ten yeast strains with mobility data in our sample (Table 3). *S. cerevisiae* strains can have high (S288c), intermediate (DBVPG6044 and Y12), or low (UWOPS05-787.3, and YPS606) Ty1 copy number, or are Ty1-less (UWOPS05-227.2 and L1374). *S. paradoxus* strains either have low copy number (CBS432 and N-44) or are Ty1-less (YPS138). All strains contain truncated Ty1 elements and Ty1-like solo LTRs, indicating that Ty1 was present in the ancestor of all strains in both species and that Ty1-less strains arose by multiple independent losses of full-length Ty1 elements, presumably by LTR-LTR recombination. Integrating genomic Ty1 content with mobility data, we observe that all restrictive strains contain full-length Ty1 elements (*S. cerevisiae*: S288c, Y12, DBVPG6044; *S. paradoxus*: CBS432, N-44), consistent with the expectation that repression of Ty1 mobility is mediated by Ty1 CNC. Also consistent with predictions of the Ty1 CNC mechanism, Ty1-less strains are permissive (*S. cerevisiae*: UWOPS05-227.2, L1374; *S. paradoxus*: YPS138). However, we also observed two permissive strains in *S. cerevisiae* that unexpectedly contained full-length Ty1 elements (UWOPS83-787.3, YPS606). These results indicate that variation in the frequency of Ty1 mobility across strains cannot be explained by a simple model whereby the presence of a full-length Ty1 element in the genome is sufficient to confer a restrictive phenotype.

### Recombination occurs among canonical Ty1 and Ty1’ subfamilies in *S. cerevisiae*

The two exceptional *S. cerevisiae* permissive strains that had full-length Ty1 elements detected in their PacBio assemblies (UWOPS05-787.3 and YPS606) displayed multiple bands with weak hybridization to the Ty1 *gag* probe by Southern blot analysis (Figure 1A). Some, but not all, of these weak Ty1 bands could be explained by cross-hybridization with Ty2 (Figure 1B). This observation suggested the possibility of divergent Ty1 sequences in these genomes such as the Ty1’ subfamily that is known to differ from the canonical Ty1 element in its *gag* region (Kim et al. 1998). To determine if the presence of a variant Ty1 subfamily could potentially explain the observation of permissive strains with full-length Ty1 elements, we extracted and aligned all full-length Ty1 elements from the PacBio assemblies of the ten *S. cerevisiae* and *S. paradoxus* strains for which we had mobility data, then clustered full-length Ty1 elements based on sequence similarity. We included the canonical Ty1-H3 tester element used in our mobility assays and used a distance-based clustering approach (Neighbor Joining) in this analysis, since our goal was to identify potential Ty1 subfamilies that could explain variation in canonical Ty1 mobility across strains, not to infer the detailed evolutionary history of Ty1 in these species.

Clustering of complete Ty1 sequences revealed a well-supported long branch separating *S. cerevisiae* elements from those in *S. paradoxus* (Figure 2A). In *S. cerevisiae*, two major clusters of Ty1 elements are observed. One cluster corresponds to the “canonical” Ty1 subfamily as defined by the presence of the Ty1-H3 tester element in this cluster (green background, Figure 2A). Two strains have full-length elements in the canonical Ty1 cluster (S288c and DBVPG6044). The other major *S. cerevisiae* cluster (found in S288c, Y12, UWOPS83-787.3, and YPS606) contains three elements previously defined as the Ty1’ subfamily in S288c by Kim *et al*. (1998) (orange background, Figure 2A). The canonical Ty1 and Ty1’ clusters are separated by a long internal branch containing multiple short branches leading to individual Ty1 elements or small groups of closely-related Ty1 elements from Y12 and S288c. Inspection of our multiple sequence alignment revealed that one of these elements (Y12_f109; single asterisk, Figure 2A) is in fact a recombinant element derived from an exchange event between canonical Ty1 and Ty1’ sequences within the *gag* region. A second recombinant in *gag* between canonical Ty1 and Ty1’ sequences was also found in our dataset (S288c_f486; double asterisk, Figure 2A), which previously was classified as a divergent Ty1’ element by Kim *et al*. (1998) (SGD: YNLCTy1-1). Sliding window analysis showed that these two recombinant elements are essentially Ty1’ elements with fragments of canonical Ty1 sequence in their *gag* regions (Sup Fig 2A, B). Full-length elements from the canonical Ty1 or Ty1’ subfamilies are not found in *S. paradoxus*, implying that both of these subfamilies are specific to *S. cerevisiae*.

**Figure 2.**
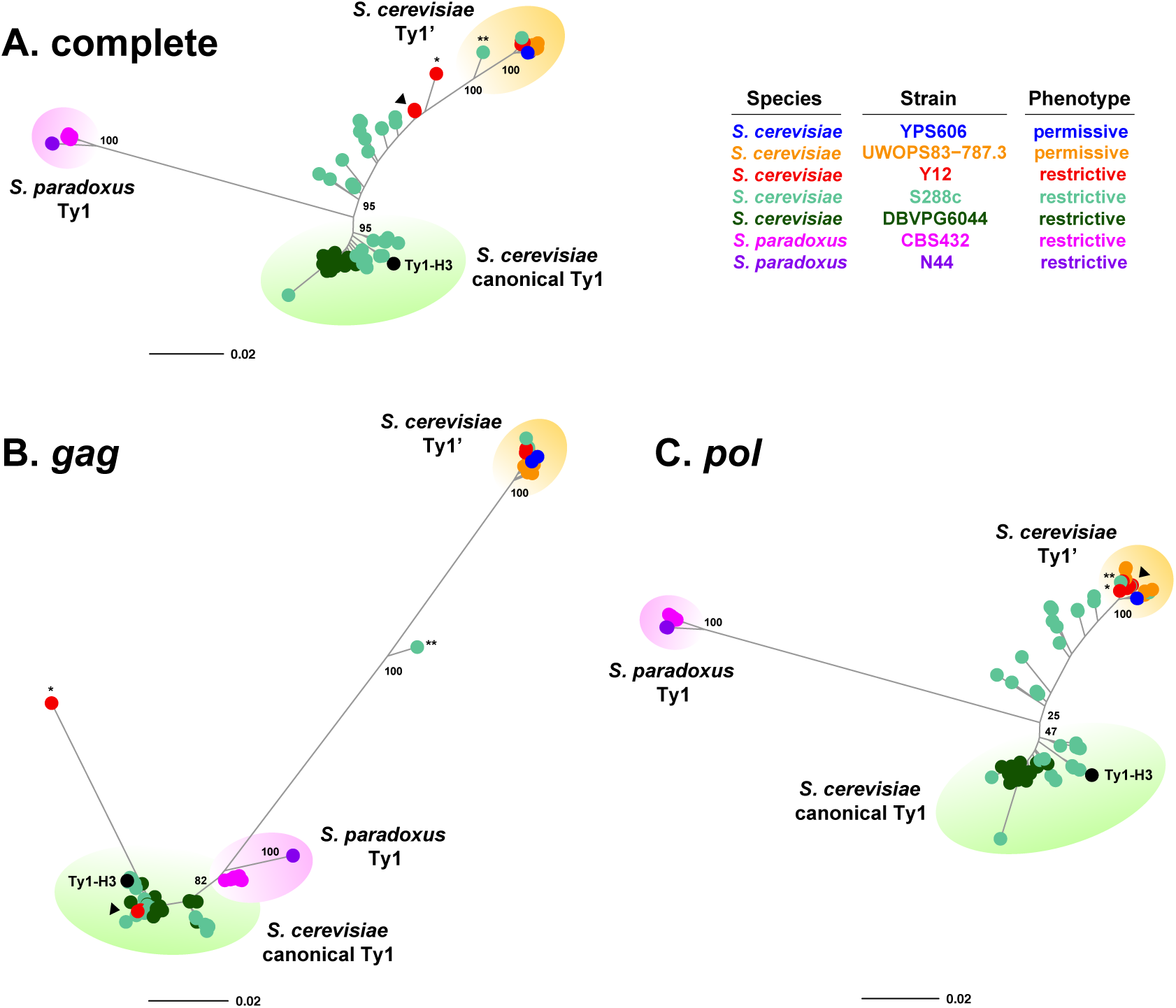
Clustering of sequences from full-length Ty1 elements in *Saccharomyces* strains with mobility phenotypes. Unrooted consensus trees for full-length elements based on (**A**) complete Ty1 sequences, (**B**) the Ty1 *gag* gene region only, and (**C**) the Ty1 *pol* gene region only. Trees we constructed using the BioNJ clustering algorithm. Numbers at key nodes represent bootstrap support based on 100 replicates. The Ty1-H3 tester element used in mobility assays is shown in black. Recombinant elements between Ty1’ and canonical Ty1 within *gag* are starred (*Y12_f109; **: S288c_f486). Seven closely related “mosaic” Y12 elements that are recombinants between canonical Ty1 and Ty1’ are labeled with an arrowhead. We note that in some cases filled circles can represent multiple identical or nearly identical sequences (e.g. seven closely related Y12 elements, arrowhead).

Because previous work shows that Ty1 *gag* and *pol* genes have different evolutionary histories in *S. cerevisiae* (Jordan and McDonald 1999b; Bleykasten-Grosshans et al. 2013), we next clustered full-length Ty1 elements on the basis of their *gag* (Figure 2B) and *pol* (Figure 2C) sequences separately. Clustering of elements based on Ty1 *gag* revealed a discordant topology relative to that from complete sequences, with no long branch separating *S. paradoxus* and *S. cerevisiae* and essentially no elements on the long internal branch separating the canonical Ty1 and Ty1’ clusters (except the recombinant S288c_f486 element noted above). In the *gag* tree, *S. paradoxus* elements unexpectedly cluster closely with *S. cerevisiae* canonical Ty1 elements, indicating a previously-unreported similarity between *S. paradoxus* Ty1 *gag* and *S. cerevisiae* canonical Ty1 *gag* (see below). Y12 and S288c elements found on the long branch between canonical Ty1 and Ty1’ clusters in the complete sequence tree (Figure 2A) cluster in the canonical group in the *gag* tree (Figure 2B), indicating that these elements all have a canonical Ty1 type *gag* gene. The recombinant Y12_f109 element noted above clusters with the canonical Ty1 group (single asterisk, Figure 2B) since the majority of its *gag* gene is canonical Ty1 but is found on a very long unique branch due to the presence of Ty1’ sequences in the 5’ part of its *gag* (Sup Fig 2A). Aside from these two elements with evidence of recombination in *gag*, all full-length Ty1 elements are found in two main groups separated by substantial sequence divergence in their *gag* region: (i) elements with a canonical Ty1 type *gag* found in *S. cerevisiae* and *S. paradoxus*, and (ii) elements with a Ty1’ type *gag* found only in *S. cerevisiae*.

Clustering of Ty1 *pol* (Figure 2C) revealed a topology similar to that of complete Ty1 sequences (Figure 2A) with two notable exceptions. First, the two elements that are recombinant in *gag* (Y12_f109 and S288c_f486) are both found within the Ty1’ cluster in their *pol* regions, consistent with these elements being predominantly Ty1’ except for parts of their *gag* genes (Sup Fig 2A, B). Second, the seven closely-related Y12 elements found on the long internal branch in the complete tree (arrowhead, Figure 2A) cluster in the Ty1’ group in the *pol* tree (arrowhead, Figure 2C). This observation, in addition to the fact that these seven Y12 elements have a canonical Ty1 *gag* (arrowhead, Figure 2B), implies that they are recombinants between canonical Ty1 and Ty1’ with an exchange event somewhere near the boundary of *gag* and *pol* (see below). Phylogenetic network analysis revealed that the remaining S288c elements that fall on the long internal branch in the *pol* tree, plus additional elements in the canonical Ty1 cluster (including the Ty1-H3 tester element), are recombinants between the canonical Ty1 and Ty1’ subfamilies within the *pol* region (Sup Fig 3). Canonical Ty1 elements from DBVPG6044, however, show no evidence of recombination in *pol* (or *gag*) and therefore best represent “pure” canonical Ty1 elements. Thus, all *S. cerevisiae* elements found on the long internal branch between canonical Ty1 and Ty1’ subfamilies in the complete sequence tree exhibit recombination between canonical Ty1 and Ty1’ within *gag*, near the boundary of the *gag* and *pol*, or within *pol*.

### *Saccharomyces* strains that restrict Ty1-H3 mobility encode full-length elements with canonical Ty1 *gag*

We next attempted to interpret variation in canonical Ty1 mobility at the strain level with genomic Ty1 content partitioned by the type of *gag* – canonical Ty1 or Ty1’ – encoded by full-length elements. The rationale for this analysis is based on p22 being encoded in the C-terminal half of *gag*, and recombination between the canonical Ty1 and Ty1’ subfamilies precluding straightforward classification at the complete element level. We classified *S. paradoxus* Ty1 elements as having canonical Ty1 type *gag* because of the close clustering with *S. cerevisiae* canonical Ty1 sequences (Figure 2B). This analysis revealed that restrictive strains from both *S. cerevisiae* and *S. paradoxus* contain one or more full-length element that encodes a canonical Ty1 type *gag* (*S. cerevisiae*: S288c, DBVPG6044 and Y12; *S. paradoxus*: CBS432 and N-44) (Table 3). Conversely, permissive strains only have full-length elements that encode a Ty1’ type *gag* (*S. cerevisiae*: UWOPS83-787.3 and YPS606) or lack full-length Ty1 elements altogether (*S. cerevisiae*: UWOPS05-227.2, L1374; *S. paradoxus*: YPS138). These results suggest that the ability to restrict mobility of a canonical Ty1-H3 tester element requires a full-length Ty1 element in the genome with sufficient sequence similarity to canonical Ty1 *gag*. Because the two restrictive strains with recombinants between canonical Ty1 and Ty1’ in *gag* (Y12 and S288c) each also have full-length elements with a complete canonical Ty1 *gag* gene, the presence of these two recombinants does not alter this general conclusion.

A confounding factor to the interpretation that a canonical Ty1 *gag* confers the restrictive phenotype is that there is substantial sequence divergence between canonical Ty1 and Ty1’ not only in *gag* but also in the LTRs and *pol* (Figure 3A). However, the impact of divergence in *gag* from these other changes between canonical Ty1 and Ty1’ can be addressed using data from the restrictive strain Y12. Full-length elements in this strain are either pure Ty1’ elements or “mosaic” Ty1 elements that have an essentially-complete canonical Ty1 *gag* but are nearly identical to Ty1’ in their LTRs and *pol* regions (Figure 3B). Remarkably, the mosaic Ty1 elements in Y12 have recombination breakpoints that coincide almost precisely with the *gag* ORF (Figure 3B), which explains the discordant placement of these elements when clustered by *gag* versus *pol* (arrowheads, Figure 2B vs. 1C). Thus, the main difference in Ty1 content between Y12 and other strains that only carry full-length Ty1’ elements (i.e. UWOPS83-787.3 and YPS606) is the presence of mosaic elements in Y12 that encode a canonical Ty1 *gag*. Given that *S. cerevisiae* strains carrying only pure Ty1’ elements have a permissive phenotype (i.e. UWOPS83-787.3 and YPS606), the restrictive phenotype in Y12 suggests that having canonical Ty1 *gag* sequences is sufficient (and that canonical LTRs and *pol* are not required) to restrict canonical Ty1-H3 mobility.

**Figure 3.**
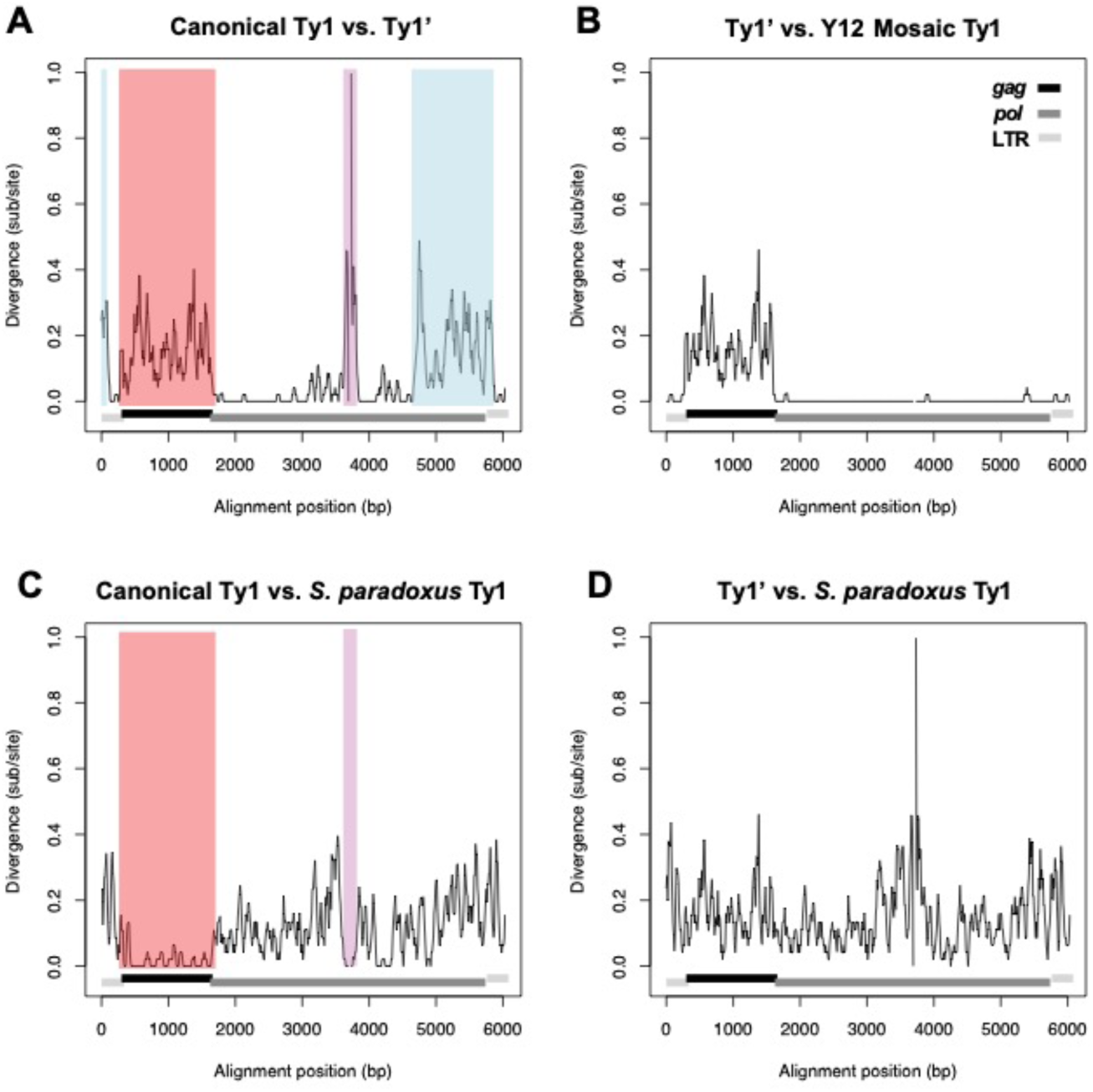
Sequence divergence between Ty1 elements in *S. cerevisiae* and *S. paradoxus*. Sliding window analysis of pairwise sequence divergence between (**A**) *S. cerevisiae* canonical Ty1 vs. *S. cerevisiae* Ty1’, (**B**) *S. cerevisiae* Ty1’ vs. *S. cerevisiae* mosaic Ty1 (found in the restrictive Y12 strain), (**C**) *S. cerevisiae* canonical Ty1 vs. *S. paradoxus* Ty1, and (**D**) *S. cerevisiae* Ty1’ vs. *S. paradoxus* Ty1. Functional regions of the Ty1 element are annotated as follows: LTRs (light grey); *gag* (black); *pol* (dark grey). Colored regions in (**A**) depict regions of high divergence between *S. cerevisiae* canonical Ty1 vs. *S. cerevisiae* Ty1’: blue: divergence in LTRs and *pol* caused by recombination between canonical Ty1 and Ty2; red: divergence in *gag* caused by recombination between canonical Ty1 and *S. paradoxus* Ty1; purple: divergence in *pol* caused by recombination between canonical Ty1 and *S. paradoxus* Ty1. Regions with high divergence between *S. cerevisiae* canonical Ty1 vs. *S. cerevisiae* Ty1’ caused by recombination with *S. paradoxus* Ty1 show low divergence between *S. cerevisiae* canonical Ty1 vs. *S. paradoxus* Ty1 in (**C**). Regions with high divergence between *S. cerevisiae* canonical Ty1 vs. *S. cerevisiae* Ty1’ caused recombination between Ty1 and Ty2 show low divergence between Ty2 and canonical Ty1 (blue, Sup Fig 4A). Elements used are: DBVPG6044_f486 (*S. cerevisiae* canonical Ty1); Y12_f208 (*S. cerevisiae* Ty1’); Y12_f429 (*S. cerevisiae* mosaic Ty1); CBS432_f139 (European *S. paradoxus* Ty1). Divergence measured in substitutions per site was calculated using a Kimura 2-parameter model in overlapping 50 bp windows with a 10 bp step size.

### Canonical *S. cerevisiae* Ty1 *gag* was recently acquired from *S. paradoxus* by horizontal transfer

Clustering of Ty1 sequences from strains with mobility data revealed an unexpected similarity between the *gag* sequences of full-length elements from the *S. cerevisiae* canonical Ty1 cluster and *S. paradoxus* (Figure 2B). Sliding window divergence analysis between canonical Ty1 and *S. paradoxus* Ty1 elements revealed that the regions of high divergence between canonical Ty1 and Ty1’ in *gag* (red, Figure 3A) and the middle part of *pol* (purple, Figure 3A) correspond exactly to regions of high sequence similarity between canonical Ty1 and *S. paradoxus* Ty1 (red and purple, Figure 3C). No such regions of high sequence similarity are observed between Ty1’ and *S. paradoxus* Ty1 (Figure 3D). These results suggest that the extreme divergence between canonical Ty1 and Ty1’ in *gag* that underlies variation in CNC phenotypes may have resulted from horizontal transfer of a *S. paradoxus* Ty1 element and recombination onto an ancestor of the *S. cerevisiae* canonical Ty1 lineage and that the Ty1’ subfamily represents the ancestral state in *S. cerevisiae*.

To further investigate this putative horizontal transfer event and the ancestral state of the Ty1 family in *S. cerevisiae*, we reconstructed the phylogenetic history of *gag* and *pol* sequences from full-length Ty1 elements using an expanded set of *S. cerevisiae* and *S. paradoxus* strains with essentially-complete PacBio assemblies. This expanded dataset includes the ten strains analyzed above, plus six *S. cerevisiae* strains with diverse origins (SK1, YPS128, UWOPS03-461.4, DBVPG6765, Sb-biocodex, Sb-unique28) and two New World *S. paradoxus* strains (UFRJ50816, UWOPS91-917.1) with publicly-available PacBio data (Khatri et al. 2017; Yue et al. 2017). Additionally, we generated and included PacBio assemblies for three *S. cerevisiae* strains isolated from ancestral oak habitats (SDO2s1, ZP568s1, ZP655.1A) to better sample Ty1 content in wild *S. cerevisiae* lineages (Almeida et al. 2015; Tilakaratna and Bensasson 2017). Ty1 sequences from PacBio assemblies of two *S. jurei* strains (NCYC3947, NCYC3962) reported in Naseeb *et al*. (2018) were used to root trees and polarize changes on the *S. cerevisiae* and *S. paradoxus* lineages. Ty elements in these genomes were detected as described above and Ty content for all strains can be found in Sup Table 1. Several strains in the expanded dataset in addition to those noted above were found to be Ty1-less and therefore are not represented in these trees: *S. cerevisiae* Wine/European (DBVPG6765, Sb-biocodex, Sb-unique28), *S. cerevisiae* Malaysian (UWOPS05-227.2, UWOPS03-461.4) and *S. paradoxus* N. America (YPS138). For this analysis, trees were generated using maximum likelihood so that ancestral states could be reconstructed for the *gag* gene. Two recombinants in *gag* noted above (S288c_f486 and Y12_f109) were excluded from this analysis since their inclusion distorted ancestral state reconstruction. Annotated trees for *gag* and *pol* with element identifiers can be found in Sup Fig 5, and Newick tree files for *gag* and *pol* can be found in Sup Files 2 and 3, respectively.

Analysis of maximum likelihood phylogenetic trees from this expanded set of strains revealed strikingly discordant histories for Ty1 *gag* (Figure 4A) and *pol* (Figure 4B). Importantly, the phylogenetic history of the *gag* gene is not compatible with the accepted species tree for these taxa (Kellis et al. 2003). In the *gag* tree, *S. cerevisiae* Ty1 sequences are found in two well-supported monophyletic groups (brown background, Figure 4A). One *S. cerevisiae gag* clade is the sister group to the ancestor of all *S. paradoxus* Ty1 *gag* sequences and contains only elements with Ty1’ type *gag*. The other *S. cerevisiae gag* clade – which includes the canonical Ty1-H3 tester element – is discordantly placed as being derived from the Old World clade of *S. paradoxus* Ty1 elements (black arrow, Figure 4A), with the closest affinity to elements from the European lineage of *S. paradoxus* represented by CBS432 (Sup Fig 5A). Importantly, all *S. cerevisiae* strains isolated from wild sources only contain elements from the Ty1’ clade, despite being sampled from diverse geographic regions around the globe: the Caribbean (UWOPS83-787.3), North America (YPS606, YPS128, SDO2s1), Asia (ZP655.1A), and Europe (ZP568s1) (Figure 4A, Sup Fig 5A). In contrast, human-associated *S. cerevisiae* strains have only canonical Ty1 elements (DBVPG6044, SK1) or both canonical Ty1 elements and Ty1’ (S288C, Y12) (Figure 4A, Sup Fig 5). The Ty1’ *gag* lineage found in human-associated strains is monophyletic, suggesting a single origin for the introduction of Ty1’ into human-associated strains of *S. cerevisiae* (grey arrow, Figure 4A). With the exception of the European *S. paradoxus* lineage unexpectedly containing sequences from the *S. cerevisiae* canonical Ty1 subfamily, the phylogeny of *S. paradoxus* Ty1 *gag* sequences follows the accepted population structure for this species (Liti et al. 2009): *S. paradoxus* Ty1 elements form an Old World clade comprised of subclades of elements from European (CBS432) and Far Eastern (N-44) lineages, plus a New World clade comprised subclades of elements from S. American (UFRJ50816) and Hawaiian (UWOPS91-917.1) lineages (Figure 4A, Sup Fig 5A).

**Figure 4.**
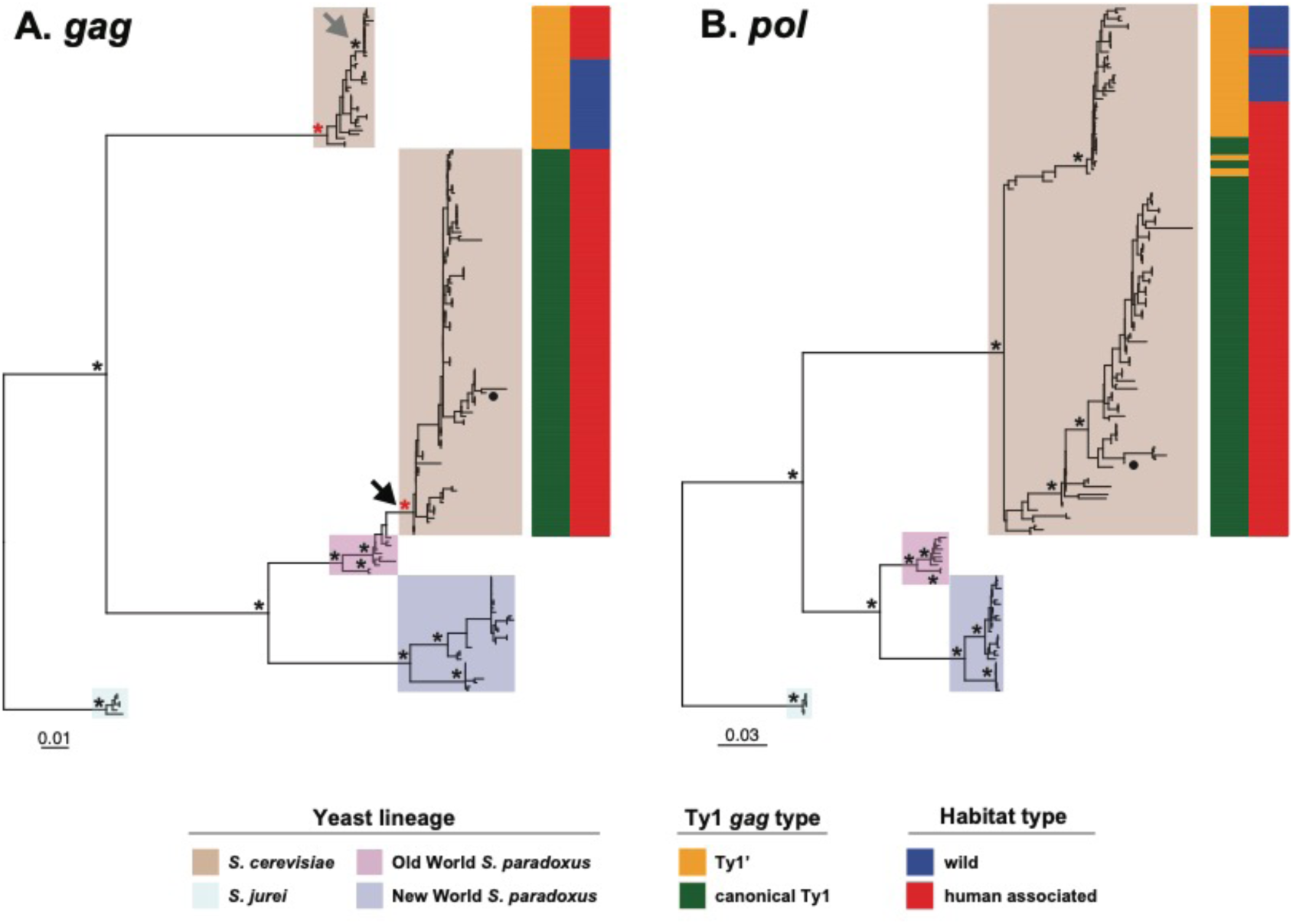
Phylogeny of *gag* and *pol* genes from full-length Ty1 elements in *S. cerevisiae* and *S. paradoxus*. Maximum likelihood phylogenies of (**A**) *gag* and (**B**) *pol* genes from full-length Ty1 elements in complete PacBio assemblies from 15 strains of *S. cerevisiae* and *S. paradoxus*, plus two strains of the outgroup species *S. jurei*. The scale bar for branch lengths is in units of substitutions per site. Asterisks represent key nodes in the phylogeny with bootstrap support >80%. The filled black circle represents the canonical Ty1-H3 element used in mobility assays. Red asterisks in (**A**) represent nodes for the ancestors of all canonical Ty1 and Ty1’ *gag* sequences, respectively, whose sequences are reconstructed in Figure 5. The black arrow in (**A**) shows evidence that the canonical Ty1 *gag* in *S. cerevisiae* is derived from an Old-World *S. paradoxus* lineage. The grey arrow in (**A**) shows evidence that elements with Ty1’ *gag* in human-associated *S. cerevisiae* have a monophyletic origin derived from a wild *S. cerevisiae* ancestor.

In contrast to the *gag* phylogeny, the *pol* tree shows species-specific clustering of Ty1 sequences for both *S. cerevisiae* and *S. paradoxus* (Figure 4B). Within *S. cerevisiae*, *pol* sequences form two major groups corresponding generally to the Ty1’ and canonical Ty1 *gag* clades (Figure 4B), but whose coherence and bootstrap support is obscured by recombination between the canonical Ty1 and Ty1’ subfamilies (as shown above for the mobility dataset). The first major *S. cerevisiae pol* clade contains all elements that have a Ty1’ *gag*, plus several other recombinant Ty1 elements that have a canonical Ty1 *gag* (e.g. the mosaic Ty1 elements from Y12). The second major *S. cerevisiae pol* clade contains elements that have a canonical Ty1 *gag* and includes the canonical Ty1-H3 tester element. Similar to *gag*, all wild strains have a Ty1’ type *pol* and all elements with a canonical Ty1 type *pol* are from human-associated strains. Divergence between the two major *S. cerevisiae* groups in *pol* is primarily caused by the regions of *pol* that underwent recombination between canonical Ty1 and either *S. paradoxus* Ty1 or Ty2, since phylogenetic analysis of a smaller region of *pol* (nucleotides 1700-3000 in Ty1-H3; Genbank: M17806) outside of the regions affected by these recombination event generates a single clade for all *S. cerevisiae* strains (Sup Fig 6, Sup File 4). As observed for *S. paradoxus* Ty1 *gag* sequences, *S. paradoxus* Ty1 *pol* sequences cluster by strain according to the accepted global biogeographic relationships for this species (Liti et al. 2009) (Figure 4B, Sup Fig 5B).

Together, these results suggest that Ty1’ is the ancestral subfamily of Ty1 in *S. cerevisiae*, as reflected by its deep divergence from *S. paradoxus* in both *gag* and *pol*, and by the unique presence of only pure Ty1’ elements (that have both Ty1’ type *gag* and *pol*) in *S. cerevisiae* strains isolated from wild habitats around the world. These results also suggest that canonical Ty1 in *S. cerevisiae* is a highly derived subfamily that acquired a complete *S. paradoxus* Ty1 *gag* sequence (as well as parts of both LTRs and *pol* from Ty2) by recombination onto a pre-existing *S. cerevisiae* Ty1’-like element, most likely in a human-associated environment. Furthermore, the placement of the *S. cerevisiae* canonical Ty1 *gag* clade within the Old World (European) *S. paradoxus* clade but sister to the New World *S. paradoxus* clade indicates this horizontal transfer event occurred in the Old World, after *S. cerevisiae* and *S. paradoxus* speciated from one another and divergence of the major worldwide *S. paradoxus* lineages had occurred.

### Divergence between canonical Ty1 and Ty1’ Gag occurs outside functionally-characterized residues

Comparison of Ty1 genomic content with mobility phenotypes above revealed that strains encoding only Ty1’ *gag* cannot strongly repress mobility of a canonical Ty1-H3 tester element (Table 1). These mobility assays imply that sequence divergence between canonical Ty1 and Ty1’ in Gag may affect the ability of Ty1’ elements to confer CNC on a canonical Ty1 element. To understand how the rate and pattern of molecular evolution acting along the branch separating the canonical Ty1 and Ty1’ subfamilies relates to potential functional divergence in *gag*, we reconstructed ancestral *gag* sequences for all canonical Ty1 elements and Ty1’ elements, respectively. Codon-based alignment of ancestral canonical Ty1 and Ty1’ *gag* sequences revealed 93 amino acid and 3 insertion/deletion substitutions across Gag, 40 of which are in the p22 region (Figure 5). Consistent with the extensive divergence in *gag* occurring along functional Ty1 lineages evolving in distinct species rather than rapid sequence evolution within species due to positive selection, we found that purifying selection was the prevailing mode of molecular evolution across both the entire *gag* gene (dN=0.125; dS=0.252; dN/dS = 0.496) and the p22 region (dN=0.128; dS=0.289; dN/dS = 0.443) (see also Kim *et al*. (1998)). Moreover, the two alternative start codons for p22 are conserved between canonical Ty1 and Ty1’ ancestors (Nishida et al. 2015; Saha et al. 2015), as are the ten CNC^R^ residues shown to provide resistance to p22 plus the seven amino acids shown to be important for Ty1 protein maturation (Tucker et al. 2015). Amino acid substitutions differentiating canonical Ty1 and Ty1’ occur at similar rates in the p22 region versus the non-p22 portions of Gag (*P*=0.907; Fisher’s Exact Test) as well as in the nine predicted helical regions relative to remaining non-helical regions of Gag (*P*=0.123; Fisher’s Exact Test). These results suggest that the Ty1’ subfamily has the capacity to code for a p22-like molecule and that potential functional divergence between canonical Ty1 and Ty1’ Gag occurs outside residues currently known to affect Ty1 protein function, maturation or resistance to p22.

**Figure 5.**
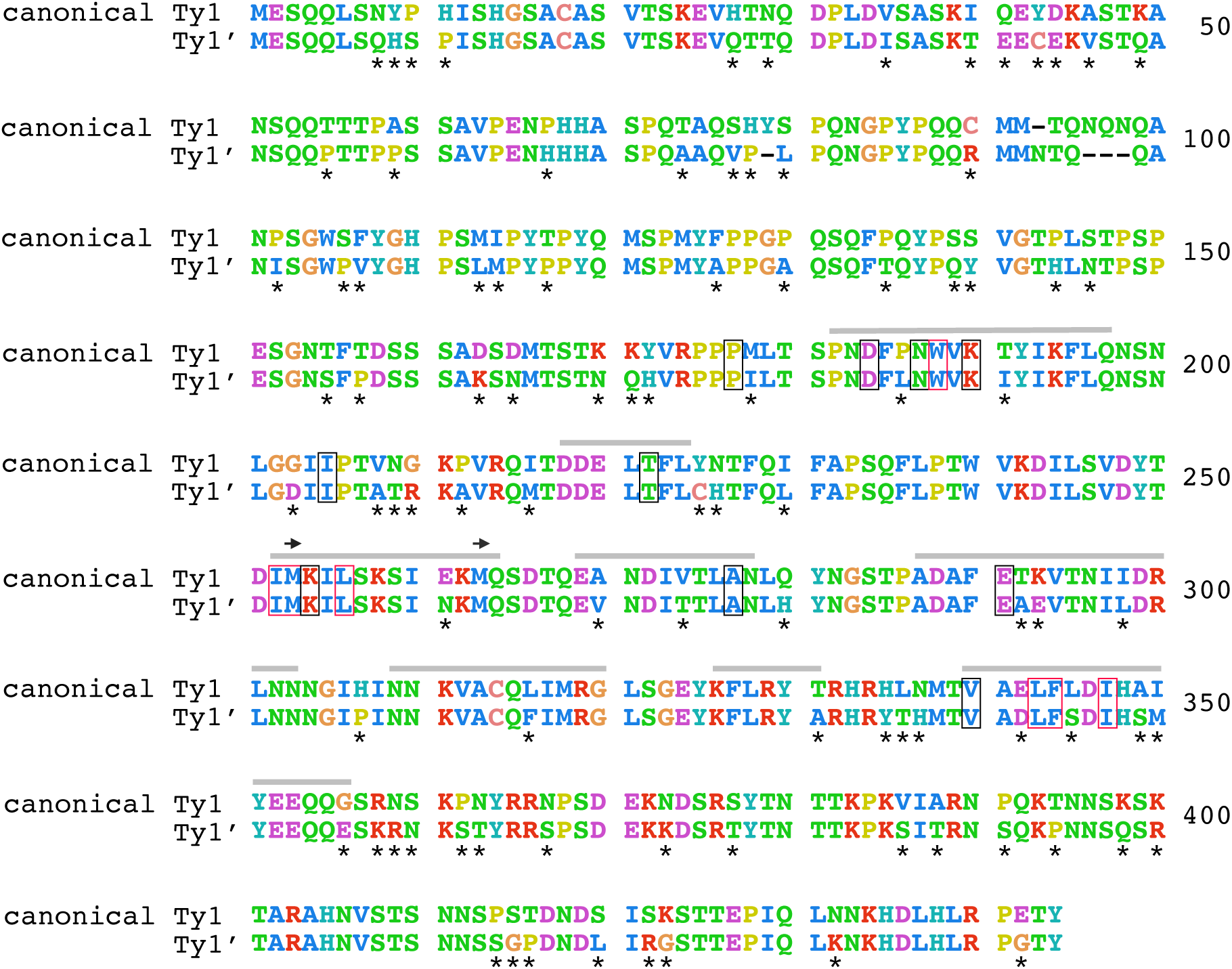
Amino acid divergence between ancestral canonical Ty1 and Ty1’ *gag* sequences. Codon-based alignment of maximum likelihood reconstructed ancestral states of all *S. cerevisiae* canonical Ty1 and all Ty1’ *gag* sequences, respectively, excluding two elements that are recombinant in *gag* (Y12_f109 and S288c_f486). Amino acids with similar properties were colorized as follows: KR (positively charged) = red; AFILMVW (hydrophobic side chains) = blue; NQST (polar uncharged side chains) = green; HY = teal; C = salmon; DE (negatively charged) = purple; P = yellow; G = orange. Amino acid substitutions between ancestral sequences are denoted by asterisks. Amino acids sites in Ty1 Gag with known function (Saha et al. 2015; Tucker et al. 2015) are annotated as follows: sites that cause resistance to p22-based CNC are shown in black boxes; sites that are important for Ty1 protein maturation are shown in red boxes; grey bars represent predicted helical regions; and alternative start codons for p22 are shown as arrows.

## Discussion

Here we combine *in vivo* transposition assays with high-resolution phylogenomics to show that variation in the ability to repress transposition of the canonical Ty1 subfamily in *S. cerevisiae* is a consequence of horizontal transfer of a *gag* gene from a closely-related yeast species, *S. paradoxus*. Our results indicate that canonical Ty1 CNC is likely to be widespread in both species and vary in strength as a function of the genomic content of specific Ty1 subfamilies. The correlation of canonical Ty1 *gag* in full-length elements with the restrictive phenotype, demonstrated most clearly by the restrictive Y12 strain, provides compelling and independent evolutionary genomic evidence to support functional studies showing that CNC is mediated by sequences within the Ty1 *gag* region (Garfinkel et al. 2003; Saha et al. 2015). Our work also reveals that, while the presence of the Ty1 family is ancestral to both species, the ability of *S. cerevisiae* strains to strongly repress canonical Ty1 CNC was acquired after speciation through horizontal transfer. Acquisition of a *S. paradoxus gag* gene by *S. cerevisiae* canonical Ty1 explains the surprising similarity between *S. cerevisiae* and *S. paradoxus* Ty1 *gag* sequences first reported here, as well as the ability of *S. paradoxus* strains with full-length Ty1 elements to restrict mobility of a heterospecific *S. cerevisiae* Ty1 tester element shown here and in previous studies (Garfinkel et al. 2003; Moore et al. 2004).

Our findings raise a number of intriguing questions about the p22-based mechanism of Ty1 CNC that can be explored in future studies. The inability of *S. cerevisiae* strains containing only Ty1’ *gag* to strongly repress canonical Ty1 implies functional divergence in the *gag* genes of these two subfamilies. While sequence analysis suggests Ty1’ is potentially capable of manifesting p22-based CNC, further work is needed to address whether this is indeed the case. If it can be shown that Ty1’ is capable of producing a functional p22 that exerts CNC on other Ty1’ elements, it will be important to evaluate which sequences in *gag*/p22 outside of currently functionally-characterized residues are responsible for this functional divergence. If Ty1’ does exert CNC *via* a p22-based mechanism, it is also possible that a Ty1’ *gag* relic present at the same locus in multiple *S. cerevisiae* strains (Bleykasten-Grosshans et al. 2013) may represent a domesticated Ty1 restriction factor similar to domesticated *gag* genes that inhibit murine or sheep retrovirus replication (Best et al. 1996; Murcia et al. 2007; Tucker and Garfinkel 2016). Functional divergence in the ability of Ty1’ to repress canonical Ty1 also raises question about potential regulatory interactions among these two subfamilies. At this point, it is clear that strains containing both canonical Ty1 and Ty1’ *gag* (e.g. native S288c, native Y12, and populated YPS606) are capable of exerting repression on canonical Ty1, and thus Ty1’ does not dominantly inhibit CNC by canonical Ty1 p22. If one assumes similar levels of Ty1*his3-AI* expression and RNA splicing in different strains, there is also some indication that permissive strains containing full-length Ty1’ elements (UWOPS83-787.3 and YPS606) have lower canonical Ty1 mobility than permissive strains that lack any full-length Ty1 elements (UWOPS05-227.2 and L-1374). This observation raises the possibility that Ty1’ can limit canonical Ty1 mobility to some degree through a CNC-like mechanism. To address these questions, future functional studies on Ty1 CNC in *S. cerevisiae* will require development and application of robust genetic techniques to characterize the transposition and CNC competence of a wider range of native Ty1 elements (including members of the Ty1’ subfamily) derived from natural isolates.

Our results also extend our understanding of the evolution of Ty1 in *S. cerevisiae* and *S. paradoxus*. Combined with previous work by Jordan and McDonald (1998; 1999b), our results suggest that the canonical Ty1 element used in most studies on Ty1 expression or function is a highly-derived element that acquired sequences from both *S. paradoxus* Ty1 and *S. cerevisiae* Ty2 in a human-associated environment. How and when these events happened remain to be determined, although the importance of homologous recombination in both events is clear. Decoding the history of the canonical Ty1 element will need to explain the somewhat paradoxical observation that this subfamily confers strong repression against itself but also apparently has high fitness, as reflected by its high copy number in strains that carry this subfamily. Understanding the history of these events may be challenging since the lack of overlap in the sequences acquired from *S. paradoxus* and Ty2 prevents their relative ordering, and the ongoing effects of recombination between canonical Ty1 and Ty1’ may obscure efforts to reconstruct the original sequences involved.

Given the common occurrence of introgression of nuclear genes from *S. paradoxus* into *S. cerevisiae* (Liti et al. 2006; Doniger et al. 2008; Strope et al. 2015; Barbosa et al. 2016; Almeida et al. 2017; Peter et al. 2018), it is also possible that multiple horizontal transfer events for Ty1 have occurred in areas where these two species coexist in nature. Thus, the presence of *S. paradoxus* sequences in canonical Ty1 *pol* (purple, Figure 3A and 2C) as well as in *gag* could either reflect independent recombination events of horizontally transferred Ty1 sequences from *S. paradoxus* or indicate that a larger fragment of *S. paradoxus* was originally transferred onto the ancestor of canonical Ty1 (with the sequence between *gag* and *pol* subsequently converted back to a Ty1’-like state). Horizontal transfer of Ty1 sequences from *S. paradoxus* into *S. cerevisiae* may have occurred by transmission of a Ty1 VLP or RNA during mating between *S. paradoxus* and *S. cerevisiae*. Alternatively, *S. paradoxus* Ty1 sequences may have entered *S. cerevisiae* by introgression of a segment of the nuclear genome carrying one or more *S. paradoxus* Ty1 elements.

Finally, our findings about the evolution of Ty1 in *S. cerevisiae* may inform aspects of the history of domestication and biogeography of this species. Recent work proposed that *S. cerevisiae* beer strains have contributions from European and Asian lineages (Fay et al. 2019). The acquisition by canonical Ty1 of sequences related to both European *S. paradoxus* Ty1 and Ty2 (which is proposed to have arisen in *S. cerevisiae* by horizontal transfer from the Asian species *S. mikatae* (Liti et al. 2005; Carr et al. 2012)) in human-associated strains supports this model. Likewise, the inference that Ty1’ is the ancestral lineage in *S. cerevisiae* and that wild strains of *S. cerevisiae* lack canonical Ty1 suggests that Ty1’ may be a useful marker for studying biogeography in this species. For example, if the “out-of-China” origin proposed for the ancestral range of wild *S. cerevisiae* is correct (Wang et al. 2012; Peter et al. 2018), our findings predict that that only the Ty1’ subfamily should be found in ancestral woodland strains of *S. cerevisiae* from China. Answers to these and other questions should be facilitated by advances in the assembly of noisy long-read datasets (Istace et al. 2017), and development of computational techniques to resolve the presence of canonical Ty1 and Ty1’ sequences from abundant short-read datasets for *S. cerevisiae* (Skelly et al. 2013; Bergstrom et al. 2014; Almeida et al. 2015; Marsit et al. 2015; Song et al. 2015; Strope et al. 2015; Barbosa et al. 2016; Barbosa et al. 2018; Duan et al. 2018; Peter et al. 2018; Fay et al. 2019; Kang et al. 2019; Ramazzotti et al. 2019).

## Materials and Methods

### Strains, plasmids, and genetic techniques

All strains used in this study are listed in Sup File 4. Strains used for Southern analysis, mobility assays and the majority of genome sequencing experiments (S288c, Y12, DBVPG6044, UWOPS83-787.3, YPS606, UWOPS05-227.2, L-1374, CBS432, N-44 and YPS138) are *MATα* haploid derivatives of *S. cerevisiae* and *S. paradoxus* strains from the SGRP (Cubillos et al. 2009). Additional monosporic wild-type *S. cerevisiae* strains used for genome sequencing (SDO2s1, ZP568s1, ZP655.1A) were reported previously (Almeida et al. 2015). For the *S. cerevisiae* SGRP strains used in mobility experiments, the *HIS3* universal gene blaster pBDG652 digested with EcoRI and SphI was used to generate *his3-Δ200hisG* deletion alleles in SGRP strains (Alani et al. 1987; Garfinkel et al. 2003). For the *S. paradoxus* strains used in mobility experiments, *KanMX* was replaced by *NatMX* by homologous recombination as previously described (Voth et al. 2003). For *S. cerevisiae* strains YPS606, UWOPS05-227.2 and L-1374, *his3-Δ200hisG* deletion strains were subsequently “populated” with canonical Ty1 elements following galactose-induced expression of pGTy1-H3 (Boeke et al. 1985), and copy number estimates were determined by Southern blot analysis as described previously (Garfinkel et al. 2003). To generate strains for mobility assay, native or populated haploid *MATα his3-Δ200hisG S. cerevisiae* strains were transformed with a Ty1*his3-AI URA3* centromere plasmid pOY1 (pBDG633) (Lee et al. 1998). Likewise native haploid *MATα NatMX S. paradoxus* strains were transformed with a Ty1*neo-AI* plasmid (pBDG954) constructed by subcloning a BstE II – Eag I fragment from pGTy1-H3PtefKAN-AI (Curcio et al. 2007) into pBDG633. Restriction endonucleases, Phusion DNA polymerase, and T4 DNA ligase were purchased from New England Biolabs (Ipswich MA). Plasmids were verified by restriction analysis and DNA sequencing. Standard yeast genetic and microbiological procedures were used in this work, including media preparation and DNA transformation (Guthrie and Fink 1991; Gietz and Schiestl 2007a; Gietz and Schiestl 2007b).

### Southern blot hybridization

A single colony from each strain was inoculated in 10 ml of YEPD medium and grown to saturation at 30°C, and total DNA was isolated as described previously (Boeke et al. 1985). Approximately 10 µg of DNA was digested with BglII and resolved on a 0.6% agarose gel for 16 hours at 30V. DNA fragments were transferred via capillary action to Hybond-N membrane (GE Healthcare, Aurora OH) and UV-crosslinked according to the supplier’s specifications (Spectroline, Westbury NY). ^32^P-labeled DNA probes containing *gag* sequences were made by randomly primed DNA synthesis with Amersham Megaprime DNA labeling System (GE Healthcare). *gag* sequences were amplified by PCR from plasmids pGTy1-H3 (Genbank: M18706; nucleotides 335 – 1496) and pGTy2-917 (Genbank: KT203716; nucleotides 333 – 1490) (Curcio et al. 1990). DNA for the Ty1 probe was amplified with primers 5’-TGGTAGCGCCTGTGCTTCGGTTAC-3’ and 5’-CATGTTTCCTCGAGTTAGTGAGCCCTGGCTGTTTCG-3’ and Phusion DNA polymerase (New England Biolabs). DNA for the Ty2 probe was generated with primers 5’-TGGTAGCGCCTATGCTTCGGTTAC-3’ and 5’-GCAATATTGTGAGCTTTTGCTGCTCTTGG-3’. Hybridization was performed at 68°C overnight in a buffer containing buffer containing 6X SSC, 5X Denhardt’s solution, 0.5% SDS, and 100 µg/µl single-strand salmon sperm DNA. Blots were washed at 68°C, twice with 2X SSC + 0.1% SDS for 30 min, followed by two washes with 1X SSC + 0.1% SDS for 15 min. Membranes were exposed to storage phosphor screens followed by scanning with Molecular Dynamics Storm PhosphorImager (GE Healthcare).

### Ty1 mobility

The frequency of Ty1*his3-AI* and Ty1*neo-AI* mobility was determined as described previously with minor modifications (Curcio and Garfinkel 1991; Garfinkel et al. 2003; Curcio et al. 2007). Briefly, a single colony from a SC-Ura plate incubated at 30°C was resuspended in 1 ml of water and 5 μl of cells was added to quadruplicate one-ml cultures of SC-Ura liquid medium. The cultures were grown for 3 days at 22°C, washed, diluted, and spread onto SC-Ura, and SC-His-Ura (for Ty1*his3-AI*) or YEPD + Geneticin (G418; 200 μg/ml for Ty1*neo-AI*) (ThermoFisher, Waltham MA) plates. To minimize flocculence, 0.25 M NaCl was used for cell washes and dilutions prior to plating (Castellon-Vogel and Menawat 1990). The frequency of Ty1 mobility is defined as the number His^+^ Ura^+^ or G418^R^ colonies divided by the number of Ura^+^ colonies per ml of culture. Standard deviations were calculated from the number Ty1 mobility events detected per 1 ml culture.

### Isolation of genomic DNA for PacBio sequencing

Genomic DNA was extracted using the Wizard Genomic DNA purification kit (Promega, Madison WI) according to the manufacturer’s instructions for yeast with minor modifications (https://dx.doi.org/10.17504/protocols.io.rved63e). A single colony of each strain was inoculated in 7 ml of YPD media and cultured for 20 - 24 hours at 30°C. Approximately 6 ml of culture for each strain was transferred to 4, 1.5 ml tubes and centrifuged at 16,162 relative centrifugal force (RCF) for 2 min at room temperature. The resulting cell pellet was resuspended in 293 µl of 50mM EDTA, pH 8.0 and incubated with 100 units of Lyticase (Sigma-Aldrich, Saint Louis MO) at 37°C for 21 hours. The resulting spheroplasts were centrifuged at 16,162 RCF for 2 min, resuspended in 300 µl of Promega Nuclei Lysis Solution, followed by adding 100 µl of Promega Protein Precipitation Solution, and incubated on ice for 5 min. Lysates were centrifuged at 16,162 RCF for 10 min. The supernatant was transferred to a 1.5 ml tube containing 300 µl of isopropanol and tubes were inverted 50 times to facilitate DNA precipitation. DNA was pelleted by centrifugation at 16,162 RCF for 10 min. The DNA pellet was washed with 300 µl of 70% ethanol, centrifuged at 16,162 RCF for 5 min and air dried for 15 min. The DNA pellet was resuspended in 50 µl of Promega DNA rehydration solution by gentle pipetting. Samples were treated with 5.25 units of RNAse A (Qiagen, Hilden Germany) at 37°C for 1 hour. Samples were incubated at 65°C for 45 min and stored at 4°C. DNA was centrifuged at 16,162 RCF for 10 min and supernatants of the four extracts were pooled, and additional purification steps were performed as follows. Qiagen DNA Hydration solution was added to bring the final volume from ~180 µl to 200 µl/strain. Three microliters of Qiagen RNAase A Solution was added to each tube followed by incubation at 37°C for 1 hour. The sample was transferred on ice and 0.5 volume of Qiagen Protein Precipitation Solution was added, followed by 2 volumes of cold 100% ethanol. Tubes were mixed by inversion 50 times and incubated on ice for 15 min. Precipitated DNA was pelleted at 16,162 RCF for 10 min and washed with 70% ethanol followed by short spin 16,162 RCF for 3 min. The supernatant was removed, and the DNA pellet was air dried for 15 min. The DNA pellet was resuspended in 110 µl of Qiagen DNA hydration Solution by gentle pipetting, followed by incubation at 37°C for 1.5 hours to fully dissolve the DNA. An additional spin at 16,162 RCF for 10 min was performed in order to pellet any impurities. Supernatants were transferred to fresh tubes and stored at 4°C or −20°C prior to making PacBio libraries.

### Genome sequencing and assembly

PacBio sequences were generated for *S. cerevisiae* using the PacBio RS II platform (Rhoads and Au 2015). For *S. cerevisiae* genomes, total DNA was purified with 1x cleaned AMPure beads (Beckman Coulter, Pasadena CA) and the quantity and quality were assessed using Nanodrop and Qubit assays. Five micrograms of purified DNA samples were sheared to approximately 20 Kb using gTubes (Covaris, Woburn MA) at 1000 x g. Sheared DNA was recovered by purification with 1:1 vol ratio of AMPure beads. Sheared DNA was treated with Exonuclease V11 (New England Biolabs), at 37°C for 15 min. The ends of the DNA were repaired by first incubating for 20 min at 37°C with damage repair mix supplied in the SMRTbell library kit (Pacific Biosciences, Menlo Park CA). This was followed by a 5-minute incubation at 25°C with end repair mix in the SMRTbell library kit. End-repaired DNA was then cleaned using 1:1 volume ratio of AMPure beads and 70% ethanol washes. End-repaired DNA was ligated to SMRTbell adapter overnight at 25°C. Ligation was terminated by incubation at 65°C for 10 min followed by exonuclease treatment for 1 hour at 37°C. The SMRTbell library was purified with 1:1 volume ratio of AMPure beads. The quantity of library and therefore the recovery was determined by Qubit assay and the average fragment size determined by Fragment Analyzer. Size-selection was performed on Sage Blue Pippin Prep using 0.75% agarose cassette and S1 marker. The size-selected SMRT bell was recovered using 1:1 volume ratio of AMPure beads and quantified by Qubit. The final average size was between 17-23 Kb. SMRTbell libraries were annealed to sequencing primer at values predetermined by the Binding Calculator (Pacific Biosciences) and a complex made with the DNA Polymerase (P6/C4 chemistry). The complexes were bound to Magbeads and this was used to set up the required number of SMRT cells for each sample. SMRT sequencing was performed using the PacBio RS II system with a movie time 360 min. Genome assembly was performed using the RS_HGAP_Assembly.3 protocol in the SMRT Analysis package version 2.3.0. Assembly statistics and quality were determined using Quast (version 4.2) (Gurevich et al. 2013) and Mummerplot (version 3.23) (Kurtz et al. 2004) relative to the UCSC sacCer3 reference assembly. Raw PacBio reads and assemblies were submitted to ENA under accession PRJEB33725.

### Annotation of Ty elements

HGAP assemblies for the genomes reported here plus a complementary set of HGAP assemblies from *S. cerevisiae*, *S. paradoxus*, and the outgroup species *S. jurei* (Yue et al. 2017; Naseeb et al. 2018) were used to identify Ty elements using a modified strategy similar to Carr *et al*. (2012). RepeatMasker (version 4.0.5; options: -e wublast -s -xsmall -nolow -no_is) (http://repeatmasker.org) was used to find all Ty fragments with similarity to a custom database of canonical Ty sequences derived from File S1 of Carr *et al*. (2012) that was updated to fix several small errors and include a version of the Tsu4 element from *S. paradoxus* (Bergman 2018) (Sup File 6). Inspection of raw RepeatMasker results identified a number of false positive matches to divergent sequences, LTR fragments from full-length or truncated elements were labeled as the incorrect family, tandem duplications of Ty elements that share a common LTR sequence that were incorrectly joined, and fusion of nearby solo LTRs fragments to full or truncated Ty elements. To fix these errors, we applied a series of automated filtering and editing operations to the raw RepeatMasker .out files: (i) false positive Ty predictions were filtered out of the RepeatMasker .out file by removing all matches to fragments with >20% divergence to the canonical Ty internal or LTR sequence; (ii) LTR fragments from full-length Ty1 and Ty2 elements were modified to match the name of the internal region found in contiguous clusters of Ty fragments with the same RepeatMasker id; (iii) tandemly-arrayed Ty elements that share an internal LTR sequence were split into distinct copies with shared LTR sequences being represented in each component copy; and (iv) flanking solo LTRs that were incorrectly joined to a complete or truncated element were split into distinct, non-overlapping copies. The final modified RepeatMasker .out file was converted to BED12 format with all fragments having the same RepeatMasker ID joined into a single BED12 record. Each Ty element in the BED12 file was categorized structurally as full-length (f, internal region present and total length >95% of canonical length), truncated (t, internal region present and total length <95% of canonical length), or solo LTRs (s, LTR present but no match to internal region). BED12 files for each strain (Sup File 7) were then used to summarize counts of all Ty element structural classes. Following Yue *et al*. (2017), counts of solo LTRs for Ty1 and Ty2 were pooled because of the similarity of their LTR sequences.

### Alignment and sequence analysis of Ty1 elements

BED12 files were used to extract fasta sequences oriented relative to the positive strand of each full-length Ty element using BEDtools getfasta (version 2.26.0; options: -name -s) (Quinlan and Hall 2010). Fasta files of full-length Ty1 sequences from all strains were concatenated together with the Ty1-H3 and Ty2 canonical elements and aligned using mafft (version 7.273) (Katoh et al. 2005) (Sup File 8). *gag* and *pol* regions were annotated based on Ty1-H3 coordinates in the resulting multiple alignment. For sliding window analysis, a subset of aligned fasta sequences was extracted and gap-only sites were removed using Seaview (version 4.7) (Gouy et al. 2010). Sequence divergence between pairs of Ty1 elements was estimated using Kimura’s 2-parameter substitution model was calculated for 50 bp windows with a 10 bp step size in R (version 3.5.2) using the spider (version 1.5) and phangorn (version 2.4) packages (Schliep 2011; Brown et al. 2012). For clustering analysis, complete sequence and region-specific alignments (for *gag* and *pol*) from the subset of strains with mobility data plus Ty1-H3 were extracted and gap-only sites were removed using Seaview (version 4.7) (Gouy et al. 2010). Aligned sequences were then clustered using the BIONJ algorithm with a Kimura 2-parameter substitution model in Seaview (version 4.7) (Gascuel 1997; Gouy et al. 2010) and resulting trees were visualized in R (version 3.5.2) using the APE package (version 5.2) (Paradis et al. 2004). Phylogenetic networks of sequences from the mobility dataset plus Ty1-H3 were generated using uncorrected P-distance and the Neighbor Net algorithm in SplitsTree 4.15.1 (Bryant and Moulton 2004; Huson and Bryant 2006). For maximum-likelihood phylogenetic analysis, region-specific alignments for the expanded dataset plus Ty1-H3 (excluding one recombinant element each from S288c and Y12) was performed using raxmlHPC-PTHREADS-AVX (version: 8.2.4; options -T 28 - x 12345 -p 12345 -b 12345 -N 100 -m GTRGAMMA) (Stamatakis 2014). Resulting phylogenetic trees were visualized in FigTree 1.4.4. Ancestral states for *gag* nucleotide sequences in the maximum likelihood tree were reconstructed using raxmlHPC-PTHREADS-AVX (version: 8.2.4; options -T 28 -f A -m GTRGAMMA) (Stamatakis 2014). Fasta sequences for the ancestors of all canonical Ty1 and Ty1’ *gag* sequences, respectively, were extracted and re-aligned using PRANK (v.170427; options: -codon -F) (Loytynoja and Goldman 2008). Codon-aligned ancestral *gag* sequences were used to estimate dN, dS, and dN/dS ratios under PAML model M0 using ETE3 (Yang 2007; Huerta-Cepas et al. 2016). Codon-based nucleotide alignments were converted to amino acids and visualized in Seaview (version 4.7) (Gouy et al. 2010).

## Acknowledgements

We thank Margaret Hughes and Xuan Liu (University of Liverpool Centre for Genomic Research) for assistance with PacBio sequencing; Joan Curcio (Wadsworth Center) for plasmid pGTy1PTefKAN-AI; Ian Donaldson and Daniela Delneri (University of Manchester) for access to the *Saccharomyces jurei* NCYC3962 assembly; Shan-Ho Tsai and Yecheng Huang (University of Georgia) for bioinformatics application support; and the Georgia Advanced Computing Resource Center for computing time. We thank members of the Bensasson, Bergman, Dyer, Garfinkel, Hall, Sweigart, White and Ye Labs at the University of Georgia for helpful suggestions throughout the project. This work was funded by the University of Georgia Research Foundation (DB, DJG and CMB), NERC NBAF award NBAF907 (DB), NIH grant R01GM095622 (DJG), and NIH grant R01GM124216 (DJG and CMB).

## Supplemental Figures

**Supplemental Figure 1. Southern blots of canonical Ty1-H3 *gag* hybridized with total DNA from *Saccharomyces* Genome Resequencing Project (SGRP) strains.** (**A**) Schematic of the Ty1 element, showing the *gag* and *pol* open reading frames (rectangles) and LTRs (arrowheads). The radiolabeled probe used for Southern blots is obtained from the *gag* gene (underlined). The location of the BglII restriction site within Ty1-H3 *pol* is shown above the schematic. Southern blot results for (**B**) *S. cerevisiae* and (**C**) *S. paradoxus*. Dots in panels (B) and (C) represent strains that were selected for Ty1 mobility assays.

**Supplemental Figure 2. Sequence divergence between Ty1’ and elements with recombination between canonical Ty1 and Ty1’ in the *gag*.** Sliding window analysis of pairwise sequence divergence between (**A**) Y12_f109 vs. Ty1’ and (**B**) S288c_f486 vs. Ty1’. The Ty1’ used in both panels is Y12_f208. Coordinates shown are relative to the multiple sequence alignment and are therefore the same for all panels. Divergence measured in substitutions per site was calculated using a Kimura 2-parameter model in overlapping 50 bp windows with a 10 bp step size.

**Supplemental Figure 3. Phylogenetic networks Ty1 sequences from *S. cerevisiae* strains with mobility phenotypes.** Neighbor-Net phylogenetic networks using uncorrected P-distances were constructed using (**A**) complete sequences, (**B**) *gag* sequences, or (**C**) *pol* sequences from full-length Ty1 elements. Nodes where incompatible partitions of sequence variation (“splits”) occur in the data because of recombination between canonical Ty1 and Ty1’ are connected by multiple edges. Bands of parallel edges would collapse to individual branches if no conflicting splits due to recombination existed in the data. Recombinant elements between Ty1’ and canonical Ty1 within *gag* are starred (*Y12_f109; **: S288c_f486).

**Supplemental Figure 4. Sequence divergence between Ty2 and Ty1 subfamilies.** Sliding window analysis of pairwise sequence divergence between (**A**) Ty2 vs. canonical Ty1, and (**B**) Ty2 vs. Ty1’. Recombination between Ty1 and Ty2 must have occurred on an ancestor of the canonical Ty1 subfamily since divergence between canonical Ty1 and Ty1’ in the LTRs and 3’ region of *pol* (blue, Figure 3) spans the same regions that have high similarity between Ty2 and canonical Ty1 (blue, Sup Fig 4A) but not between Ty2 and Ty1’ (Sup Fig 4B). Identifiers for elements shown are: RepBase TY2#LTR/Copia (Ty2), DBVPG6044_f486 (canonical Ty1); Y12_f208 (Ty1’). Coordinates shown are relative to the multiple sequence alignment and are therefore the same for all panels. Divergence measured in substitutions per site was calculated using a Kimura 2-parameter model in overlapping 50 bp windows with a 10 bp step size.

**Supplemental Figure 5. Strain-labelled phylogeny of *gag* and *pol* genes from full-length Ty1 elements in *S. cerevisiae* and *S. paradoxus*.** Maximum likelihood phylogenies of (**A**) *gag* and (**B**) *pol* genes from full-length Ty1 elements in complete PacBio assemblies from 15 strains of *S. cerevisiae* and *S. paradoxus*, plus two strains of the outgroup species *S. jurei*. The scale bar for branch lengths is in units of substitutions per site. Nodes labelled by asterisks in Fig 3 are shown with bootstrap support values. Ty1 element identifiers are a composite of strain name followed by a strain-specific unique numerical identifier prefixed by “f” indicating that it is a full-length element. Tree files in Newick format for *gag* and *pol* can be found in Sup Files 2 and 3, respectively. Aligned fasta sequences for all full-length Ty1 elements can be found in Sup File 8.

**Supplemental Figure 6. Strain-labelled phylogeny of non-recombinant *pol* region from full-length Ty1 elements in *S. cerevisiae* and *S. paradoxus*.** Maximum likelihood phylogenies of region of *pol* gene outside regions of recombination (nucleotides 1700-3000 in M18706) between canonical Ty1 and either *S. paradoxus* Ty1 or Ty2 from full-length Ty1 elements in complete PacBio assemblies from 15 strains of *S. cerevisiae* and *S. paradoxus*, plus two strains of the outgroup species *S. jurei*. The scale bar for branch lengths is in units of substitutions per site. Bootstrap support is shown for nodes with values >95%. Ty1 element identifiers are a composite of strain name followed by a strain-specific unique numerical identifier prefixed by “f” indicating that it is a full-length element. Tree file in Newick format for the non-recombinant region of *pol* can be found in Sup File 4. Aligned fasta sequences for all full-length Ty1 elements can be found in Sup File 8.

## Supplemental Files

**Supplemental File 1. Assembly statistics and Ty content in PacBio assemblies of *Saccharomyces* species.** Strains from the current work are labeled Czaja; strains from published assemblies are labeled by the last name of the first author of the respective paper (Khatri et al. 2017; Yue et al. 2017; Naseeb et al. 2018). Counts of Ty elements are based on structural classification of fragments from the same RepeatMasker annotation: full-length (internal region present and total length >95% of canonical length), truncated (internal region present and total length <95% of canonical length), or solo LTRs (LTR present but no match to internal region). Following Yue *et al*. (2017), counts of solo LTRs for Ty1 and Ty2 were pooled because of the similarity of their LTR sequences.

**Supplemental File 2. Maximum likelihood tree for the complete Ty1 *gag* region.** Newick-formatted maximum likelihood tree file based on *gag* sequences of full-length Ty1 elements in the expanded dataset plus the canonical Ty1-H3 element (M18706). Node labels represent bootstrap support based on 100 replicates and branch lengths are in substitutions per site.

**Supplemental File 3. Maximum likelihood tree for the complete Ty1 *pol* region.** Newick-formatted maximum likelihood tree file based on *pol* sequences of full-length Ty1 elements in the expanded dataset plus the canonical Ty1-H3 element (M18706). Node labels represent bootstrap support based on 100 replicates and branch lengths are in substitutions per site.

**Supplemental File 4. Maximum likelihood tree for the non-recombinant region of Ty1 *pol*.** Newick-formatted maximum likelihood tree file based on *pol* sequences corresponding to nucleotides 1700-3000 of M18706 of full-length Ty1 elements in the expanded dataset plus the canonical Ty1-H3 element (M18706). Node labels represent bootstrap support based on 100 replicates and branch lengths are in substitutions per site.

**Supplemental File 5. List of *S. cerevisiae* and *S. paradoxus* strains used in this study.** Original strains used were initially reported in Cubillos *et al*. (2009) and Almeida *et al*. (2015). Identifiers from the National Collection of Yeast Cultures are provided for original strains from Cubillos *et al*. (2009). Identifiers from the David Garfinkel lab collection are provided for modified strains carrying selectable markers or Ty1 mobility plasmids.

**Supplemental File 6. Database of Ty element query sequences.** Fasta file of LTR and internal sequences for Ty1-Ty5 from *S. cerevisiae* and Ty3p and Tsu4 from *S. paradoxus* used for RepeatMasker-based annotation of Ty elements in yeast genomes.

**Supplemental File 7. BED file of Ty element coordinates.** Strain-specific BED12 files of Ty elements for all strains in the expanded dataset. Each Ty element in the BED12 file was categorized structurally with a prefix as full-length (f, internal region present and total length >95% of canonical length), truncated (t, internal region present and total length <95% of canonical length), or solo LTRs (s, LTR present but no match to internal region). Sequence IDs correspond to International Nucleotide Sequence Database Collaboration records except for *S. jurei* NCYC3962 which uses IDs from an assembly provided by personal communication from the Daniela Delneri lab (University of Manchester).

**Supplemental File 8. Full-length Ty1 element nucleotide sequences.** Multiple sequence alignment of full-length Ty1 elements in the expanded dataset plus the canonical Ty1-H3 element (M18706). Terminal deletions exist in some sequences since Ty1 elements are classified as being full-length on the basis of having partial or complete sequences for both LTR sequences plus an internal region sequence that is >95% the length of internal region for the query Ty1 element.

## References

Ahn HW, Tucker JM, Arribere JA, Garfinkel DJ. 2017. Ribosome biogenesis modulates Ty1 copy number control in Saccharomyces cerevisiae. Genetics 207:1441–1456.

Alani E, Cao L, Kleckner N. 1987. A method for gene disruption that allows repeated use of URA3 selection in the construction of multiply disrupted yeast strains. Genetics 116:541–545.

Almeida P, Barbosa R, Bensasson D, Goncalves P, Sampaio JP. 2017. Adaptive divergence in wine yeasts and their wild relatives suggests a prominent role for introgressions and rapid evolution at noncoding sites. Mol. Ecol. 26:2167–2182.

Almeida P, Barbosa R, Zalar P, Imanishi Y, Shimizu K, Turchetti B, Legras J-L, Serra M, Dequin S, Couloux A, et al. 2015. A population genomics insight into the Mediterranean origins of wine yeast domestication. Mol Ecol 24:5412–5427.

Atwood A, Choi J, Levin HL. 1998. The application of a homologous recombination assay revealed amino acid residues in an LTR-retrotransposon that were critical for integration. J. Virol. 72:1324–1333.

Barbosa R, Almeida P, Safar SVB, Santos RO, Morais PB, Nielly-Thibault L, Leducq J-B, Landry CR, Goncalves P, Rosa CA, et al. 2016. Evidence of natural hybridization in Brazilian wild lineages of Saccharomyces cerevisiae. Genome Biol Evol 8:317–329.

Barbosa R, Pontes A, Santos RO, Montandon GG, de Ponzzes-Gomes CM, Morais PB, Gonçalves P, Rosa CA, Sampaio JP. 2018. Multiple Rounds of Artificial Selection Promote Microbe Secondary Domestication—The Case of Cachaça Yeasts. Genome Biol Evol 10:1939–1955.

Bergman CM. 2018. Horizontal transfer and proliferation of Tsu4 in Saccharomyces paradoxus. Mobile DNA 9:18.

Bergstrom A, Simpson JT, Salinas F, Barre B, Parts L, Zia A, Ba N, N A, Moses AM, Louis EJ, et al. 2014. A high-definition view of functional genetic variation from natural yeast genomes. Mol Biol Evol 31:872–888.

Best S, Le Tissier P, Towers G, Stoye JP. 1996. Positional cloning of the mouse retrovirus restriction gene Fv1. Nature 382:826–829.

Bleykasten-Grosshans C, Friedrich A, Schacherer J. 2013. Genome-wide analysis of intraspecific transposon diversity in yeast. BMC Genomics 14:399.

Boeke JD, Eichinger D, Castrillon D, Fink GR. 1988. The Saccharomyces cerevisiae genome contains functional and nonfunctional copies of transposon Ty1. Mol Cell Biol 8:1432–1442.

Boeke JD, Garfinkel DJ, Styles CA, Fink GR. 1985. Ty elements transpose through an RNA intermediate. Cell 40:491–500.

Bridier-Nahmias A, Tchalikian-Cosson A, Baller JA, Menouni R, Fayol H, Flores A, Saïb A, Werner M, Voytas DF, Lesage P. 2015. Retrotransposons. An RNA polymerase III subunit determines sites of retrotransposon integration. Science 348:585–588.

Brown SDJ, Collins RA, Boyer S, Lefort M-C, Malumbres-Olarte J, Vink CJ, Cruickshank RH. 2012. Spider: An R package for the analysis of species identity and evolution, with particular reference to DNA barcoding. Molecular Ecology Resources 12:562–565.

Bryant D, Moulton V. 2004. Neighbor-Net: An Agglomerative Method for the Construction of Phylogenetic Networks. Mol Biol Evol 21:255–265.

Carr M, Bensasson D, Bergman CM. 2012. Evolutionary genomics of transposable elements in Saccharomyces cerevisiae. PLoS ONE 7:e50978.

Castellon-Vogel MA, Menawat AS. 1990. A method to disperse aggregates of a flocculent yeast for photometric analysis. Biotechnol. Prog. 6:135–141.

Chenais B, Caruso A, Hiard S, Casse N. 2012. The impact of transposable elements on eukaryotic genomes: from genome size increase to genetic adaptation to stressful environments. Gene 509:7–15.

Cheung S, Ma L, Chan PHW, Hu H-L, Mayor T, Chen H-T, Measday V. 2016. Ty1-Integrase interacts with RNA Polymerase III specific subcomplexes to promote insertion of Ty1 elements upstream of Pol III-transcribed genes. J. Biol. Chem.

Cubillos FA, Louis EJ, Liti G. 2009. Generation of a large set of genetically tractable haploid and diploid Saccharomyces strains. FEMS Yeast Research 9:1217–1225.

Curcio MJ, Garfinkel DJ. 1991. Single-step selection for Ty1 element retrotransposition. Proc. Natl. Acad. Sci. U.S.A. 88:936–940.

Curcio MJ, Hedge A-M, Boeke JD, Garfinkel DJ. 1990. Ty RNA levels determine the spectrum of retrotransposition events that activate gene expression in Saccharomyces cerevisiae. Molec. Gen. Genet. 220:213–221.

Curcio MJ, Kenny AE, Moore S, Garfinkel DJ, Weintraub M, Gamache ER, Scholes DT. 2007. S-phase checkpoint pathways stimulate the mobility of the retrovirus-like transposon Ty1. Mol. Cell. Biol. 27:8874–8885.

Curcio MJ, Lutz S, Lesage P. 2015. The Ty1 LTR-retrotransposon of budding yeast, Saccharomyces cerevisiae. Microbiol Spectr 3:1–35.

Doniger SW, Kim HS, Swain D, Corcuera D, Williams M, Yang S-P, Fay JC. 2008. A catalog of neutral and deleterious polymorphism in yeast. PLoS Genet. 4:e1000183.

Drinnenberg IA, Weinberg DE, Xie KT, Mower JP, Wolfe KH, Fink GR, Bartel DP. 2009. RNAi in budding yeast. Science 326:544–550.

Duan S-F, Han P-J, Wang Q-M, Liu W-Q, Shi J-Y, Li K, Zhang X-L, Bai F-Y. 2018. The origin and adaptive evolution of domesticated populations of yeast from Far East Asia. Nature Communications 9:2690.

Fay JC, Liu P, Ong GT, Dunham MJ, Cromie GA, Jeffery EW, Ludlow CL, Dudley AM. 2019. A polyploid admixed origin of beer yeasts derived from European and Asian wine populations. PLOS Biology 17:e3000147.

Friedli M, Trono D. 2015. The developmental control of transposable elements and the evolution of higher species. Annu. Rev. Cell Dev. Biol. 31:429–451.

Garfinkel DJ. 2005. Genome evolution mediated by Ty elements in Saccharomyces. Cytogenet Genome Res 110:63–69.

Garfinkel DJ, Nyswaner K, Wang J, Cho J-Y. 2003. Post-transcriptional cosuppression of Ty1 retrotransposition. Genetics 165:83–99.

Garfinkel DJ, Tucker JM, Saha A, Nishida Y, Pachulska-Wieczorek K, Błaszczyk L, Purzycka KJ. 2016. A self-encoded capsid derivative restricts Ty1 retrotransposition in Saccharomyces. Curr. Genet. 62:321–329.

Gascuel O. 1997. BIONJ: an improved version of the NJ algorithm based on a simple model of sequence data. Mol. Biol. Evol. 14:685–695.

Gietz RD, Schiestl RH. 2007a. High-efficiency yeast transformation using the LiAc/SS carrier DNA/PEG method. Nat Protoc 2:31–34.

Gietz RD, Schiestl RH. 2007b. Quick and easy yeast transformation using the LiAc/SS carrier DNA/PEG method. Nat Protoc 2:35–37.

Goffeau A, Barrell BG, Bussey H, Davis RW, Dujon B, Feldmann H, Galibert F, Hoheisel JD, Jacq C, Johnston M, et al. 1996. Life with 6000 genes. Science 274:546, 563–7.

Goodier JL. 2016. Restricting retrotransposons: a review. Mob DNA 7:16.

Gouy M, Guindon S, Gascuel O. 2010. SeaView version 4: A multiplatform graphical user interface for sequence alignment and phylogenetic tree building. Mol Biol Evol 27:221–224.

Gurevich A, Saveliev V, Vyahhi N, Tesler G. 2013. QUAST: quality assessment tool for genome assemblies. Bioinformatics 29:1072–1075.

Guthrie C, Fink GR eds. 1991. Guide to yeast genetics and molecular biology. Spi edition. San Diego, Calif.: Academic Press

Huerta-Cepas J, Serra F, Bork P. 2016. ETE 3: Reconstruction, analysis, and visualization of phylogenomic data. Mol Biol Evol 33:1635–1638.

Huson DH, Bryant D. 2006. Application of phylogenetic networks in evolutionary studies. Mol. Biol. Evol. 23:254–267.

Istace B, Friedrich A, d’Agata L, Faye S, Payen E, Beluche O, Caradec C, Davidas S, Cruaud C, Liti G, et al. 2017. De novo assembly and population genomic survey of natural yeast isolates with the Oxford Nanopore MinION sequencer. Gigascience 6:1–13.

Jordan IK, McDonald JF. 1998. Evidence for the role of recombination in the regulatory evolution of Saccharomyces cerevisiae Ty elements. Journal of Molecular Evolution 47:14–20.

Jordan IK, McDonald JF. 1999a. Tempo and mode of Ty element evolution in Saccharomyces cerevisiae. Genetics 151:1341–51.

Jordan IK, McDonald JF. 1999b. Phylogenetic perspective reveals abundant Ty1/Ty2 hybrid elements in the Saccharomyces cerevisiae genome. Mol. Biol. Evol. 16:419–422.

Kang K, Bergdahl B, Machado D, Dato L, Han T-L, Li J, Villas-Boas S, Herrgård MJ, Förster J, Panagiotou G. 2019. Linking genetic, metabolic, and phenotypic diversity among Saccharomyces cerevisiae strains using multi-omics associations. Gigascience 8.

Katoh K, Kuma K, Toh H, Miyata T. 2005. MAFFT version 5: improvement in accuracy of multiple sequence alignment. Nucleic Acids Res 33:511–518.

Kellis M, Patterson N, Endrizzi M, Birren B, Lander ES. 2003. Sequencing and comparison of yeast species to identify genes and regulatory elements. Nature 423:241–54.

Khatri I, Tomar R, Ganesan K, Prasad GS, Subramanian S. 2017. Complete genome sequence and comparative genomics of the probiotic yeast Saccharomyces boulardii. Sci Rep 7:371.

Kim JM, Vanguri S, Boeke JD, Gabriel A, Voytas DF. 1998. Transposable elements and genome organization: a comprehensive survey of retrotransposons revealed by the complete Saccharomyces cerevisiae genome sequence. Genome Res. 8:464–478.

Kurtz S, Phillippy A, Delcher AL, Smoot M, Shumway M, Antonescu C, Salzberg SL. 2004. Versatile and open software for comparing large genomes. Genome Biol. 5:R12.

Lee BS, Lichtenstein CP, Faiola B, Rinckel LA, Wysock W, Curcio MJ, Garfinkel DJ. 1998. Posttranslational inhibition of Ty1 retrotransposition by nucleotide excision repair/transcription factor TFIIH subunits Ssl2p and Rad3p. Genetics 148:1743–1761.

Liti G, Barton DBH, Louis EJ. 2006. Sequence diversity, reproductive isolation and species concepts in Saccharomyces. Genetics 174:839–850.

Liti G, Carter DM, Moses AM, Warringer J, Parts L, James SA, Davey RP, Roberts IN, Burt A, Koufopanou V, et al. 2009. Population genomics of domestic and wild yeasts. Nature 458:337–41.

Liti G, Peruffo A, James SA, Roberts IN, Louis EJ. 2005. Inferences of evolutionary relationships from a population survey of LTR-retrotransposons and telomeric-associated sequences in the Saccharomyces sensu stricto complex. Yeast 22:177–192.

Loytynoja A, Goldman N. 2008. Phylogeny-aware gap placement prevents errors in sequence alignment and evolutionary analysis. Science 320:1632–1635.

Marsit S, Mena A, Bigey F, Sauvage F-X, Couloux A, Guy J, Legras J-L, Barrio E, Dequin S, Galeote V. 2015. Evolutionary advantage conferred by an eukaryote-to-eukaryote gene transfer event in wine yeasts. Mol Biol Evol 32:1695–1707.

Menconi G, Battaglia G, Grossi R, Pisanti N, Marangoni R. 2013. Mobilomics in Saccharomyces cerevisiae strains. BMC Bioinformatics 14:102.

Mita P, Boeke JD. 2016. How retrotransposons shape genome regulation. Curr. Opin. Genet. Dev. 37:90–100.

Moore SP, Liti G, Stefanisko KM, Nyswaner KM, Chang C, Louis EJ, Garfinkel DJ. 2004. Analysis of a Ty1-less variant of Saccharomyces paradoxus: the gain and loss of Ty1 elements. Yeast 21:649–60.

Murcia PR, Arnaud F, Palmarini M. 2007. The transdominant endogenous retrovirus enJS56A1 associates with and blocks intracellular trafficking of Jaagsiekte sheep retrovirus Gag. J. Virol. 81:1762–1772.

Naseeb S, Alsammar H, Burgis T, Donaldson I, Knyazev N, Knight C, Delneri D. 2018. Whole genome sequencing, de novo assembly and phenotypic profiling for the new budding yeast species Saccharomyces jurei. G3 8:2967–2977.

Nelson MG, Linheiro RS, Bergman CM. 2017. McClintock: an integrated pipeline for detecting transposable element insertions in whole-genome shotgun sequencing data. G3 7:2749–2762.

Neuveglise C, Feldmann H, Bon E, Gaillardin C, Casaregola S. 2002. Genomic evolution of the long terminal repeat retrotransposons in hemiascomycetous yeasts. Genome Research 12:930–43.

Nishida Y, Pachulska-Wieczorek K, Błaszczyk L, Saha A, Gumna J, Garfinkel DJ, Purzycka KJ. 2015. Ty1 retrovirus-like element Gag contains overlapping restriction factor and nucleic acid chaperone functions. Nucleic Acids Res. 43:7414–7431.

Paradis E, Claude J, Strimmer K. 2004. APE: Analyses of phylogenetics and evolution in R language. Bioinformatics 20:289–290.

Peter J, Chiara MD, Friedrich A, Yue J-X, Pflieger D, Bergstrom A, Sigwalt A, Barre B, Freel K, Llored A, et al. 2018. Genome evolution across 1,011 Saccharomyces cerevisiae isolates. Nature 556:339–344.

Promislow DE, Jordan IK, McDonald JF. 1999. Genomic demography: a life-history analysis of transposable element evolution. Proc Biol Sci 266:1555–60.

Quinlan AR, Hall IM. 2010. BEDTools: a flexible suite of utilities for comparing genomic features. Bioinformatics 26:841–842.

Ramazzotti M, Stefanini I, Paola MD, Filippo CD, Rizzetto L, Berná L, Dapporto L, Rivero D, Tocci N, Weil T, et al. 2019. Population genomics reveals evolution and variation of Saccharomyces cerevisiae in the human and insects gut. Environmental Microbiology 21:50–71.

Rhoads A, Au KF. 2015. PacBio Sequencing and Its Applications. Genomics, Proteomics & Bioinformatics 13:278–289.

Saha A, Mitchell JA, Nishida Y, Hildreth JE, Ariberre JA, Gilbert WV, Garfinkel DJ. 2015. A trans-dominant form of Gag restricts Ty1 retrotransposition and mediates copy number control. J. Virol. 89:3922–3938.

Schliep KP. 2011. phangorn: phylogenetic analysis in R. Bioinformatics 27:592–593.

Sharon G, Burkett TJ, Garfinkel DJ. 1994. Efficient homologous recombination of Ty1 element cDNA when integration is blocked. Mol. Cell. Biol. 14:6540–6551.

Skelly DA, Merrihew GE, Riffle M, Connelly CF, Kerr EO, Johansson M, Jaschob D, Graczyk B, Shulman NJ, Wakefield J, et al. 2013. Integrative phenomics reveals insight into the structure of phenotypic diversity in budding yeast. Genome Res. 23:1496–1504.

Song G, Dickins BJA, Demeter J, Engel S, Gallagher J, Choe K, Dunn B, Snyder M, Cherry JM.2015. AGAPE (Automated Genome Analysis PipelinE) for pan-genome analysis of Saccharomyces cerevisiae. PLoS ONE 10:e0120671.

Stamatakis A. 2014. RAxML version 8: a tool for phylogenetic analysis and post-analysis of large phylogenies. Bioinformatics 30:1312–1313.

Strope PK, Skelly DA, Kozmin SG, Mahadevan G, Stone EA, Magwene PM, Dietrich FS, McCusker JH. 2015. The 100-genomes strains, an S. cerevisiae resource that illuminates its natural phenotypic and genotypic variation and emergence as an opportunistic pathogen. Genome Res. 25:762–774.

Tilakaratna V, Bensasson D. 2017. Habitat predicts levels of genetic admixture in Saccharomyces cerevisiae. G3 7:2919–2929.

Tucker JM, Garfinkel DJ. 2016. Ty1 escapes restriction by the self-encoded factor p22 through mutations in capsid. Mob Genet Elements 6:e1154639.

Tucker JM, Larango ME, Wachsmuth LP, Kannan N, Garfinkel DJ. 2015. The Ty1 Retrotransposon Restriction Factor p22 Targets Gag. PLOS Genet 11:e1005571.

Voth WP, Jiang YW, Stillman DJ. 2003. New “marker swap” plasmids for converting selectable markers on budding yeast gene disruptions and plasmids. Yeast 20:985–993.

Wang Q-M, Liu W-Q, Liti G, Wang S-A, Bai F-Y. 2012. Surprisingly diverged populations of Saccharomyces cerevisiae in natural environments remote from human activity. Mol. Ecol. 21:5404–5417.

Wilke CM, Maimer E, Adams J. 1992. The population biology and evolutionary significance of Ty elements in Saccharomyces cerevisiae. Genetica 86:155–73.

Yang Z. 2007. PAML 4: Phylogenetic Analysis by Maximum Likelihood. Mol Biol Evol 24:1586–1591.

Yue J-X, Li J, Aigrain L, Hallin J, Persson K, Oliver K, Bergstrom A, Coupland P, Warringer J, Lagomarsino MC, et al. 2017. Contrasting evolutionary genome dynamics between domesticated and wild yeasts. Nat Genet 49:913–924.

