## Supplementary figures and images for "Evolution of Ty1 copy number control in yeast by horizontal transfer of a *gag* gene"

### Supplemental Figure 1

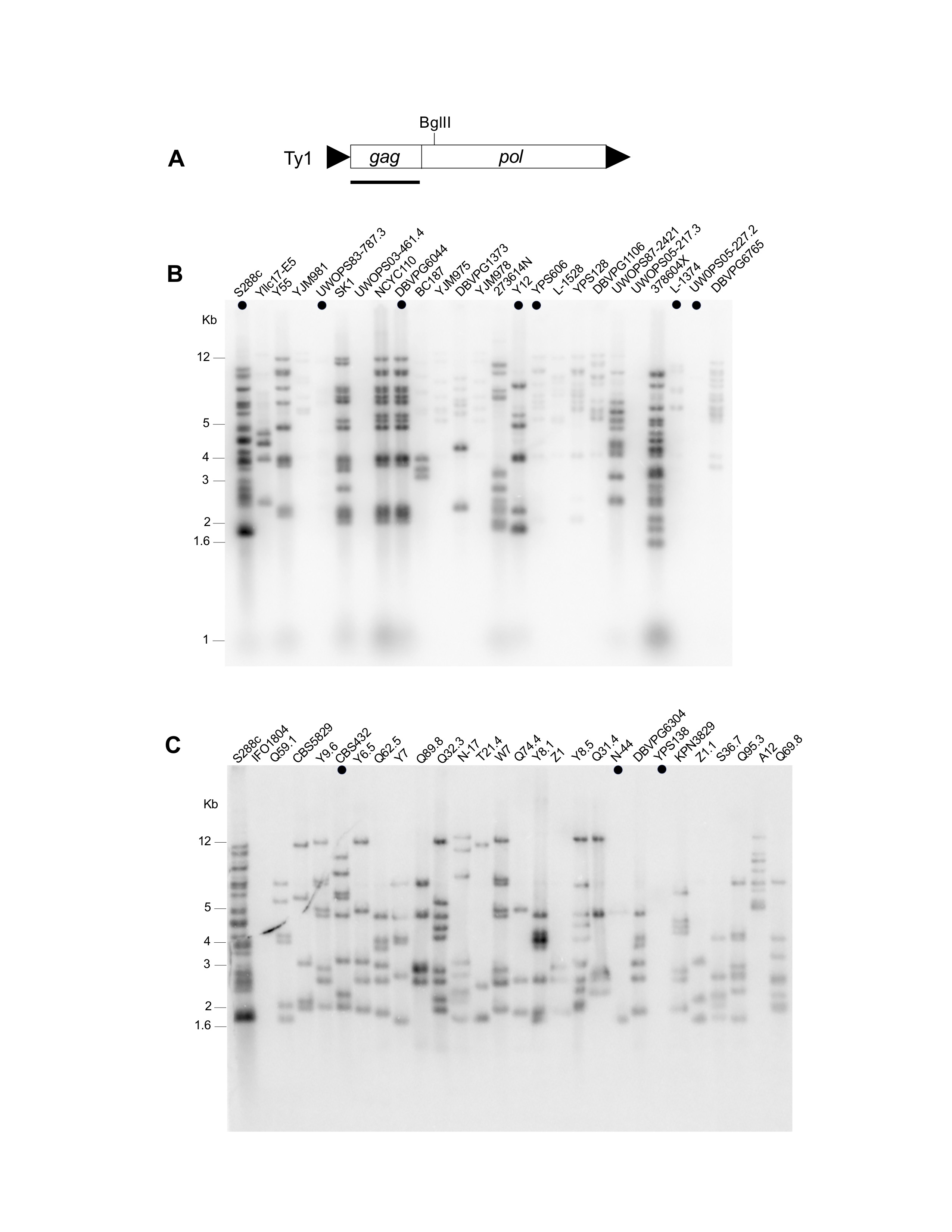

### Supplemental Figure 2

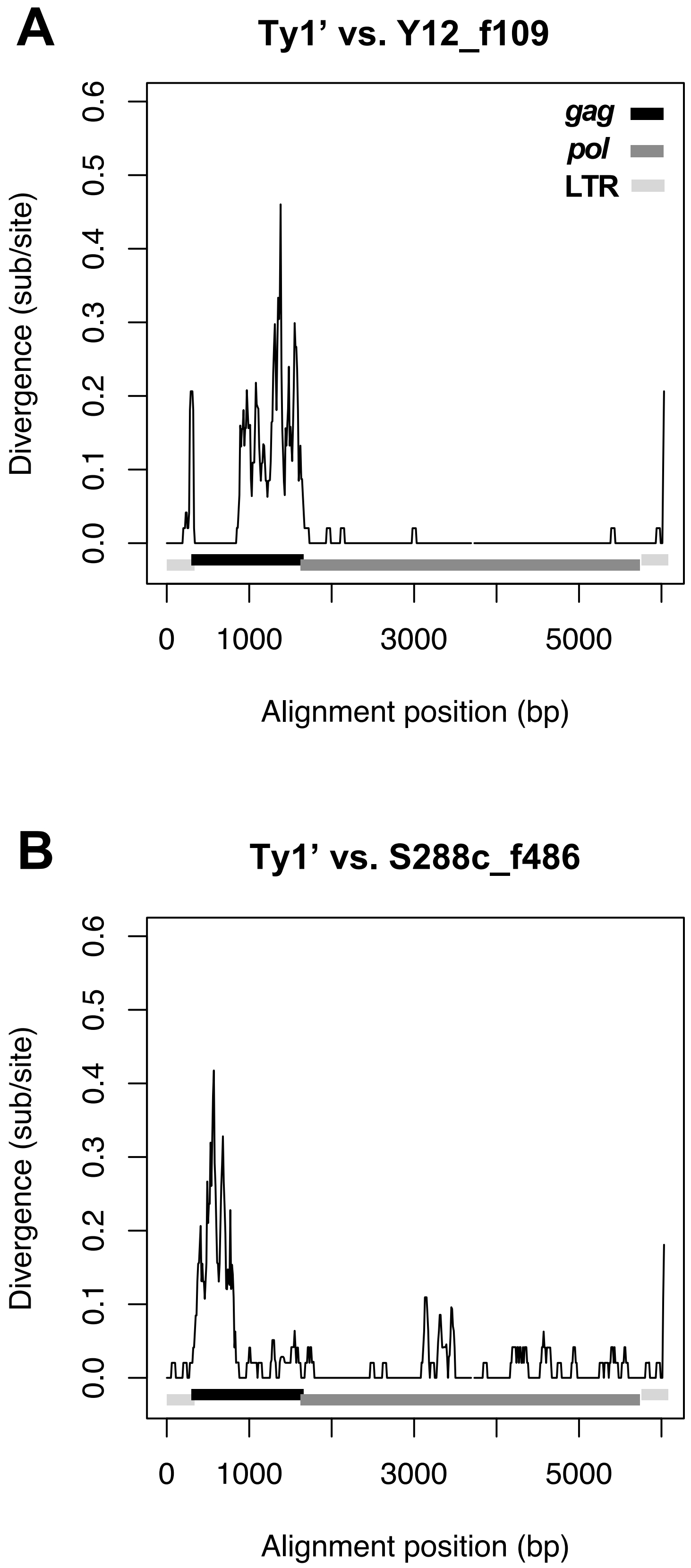

### Supplemental Figure 3

#### A. complete

#### B. *gag*

#### C. *pol*

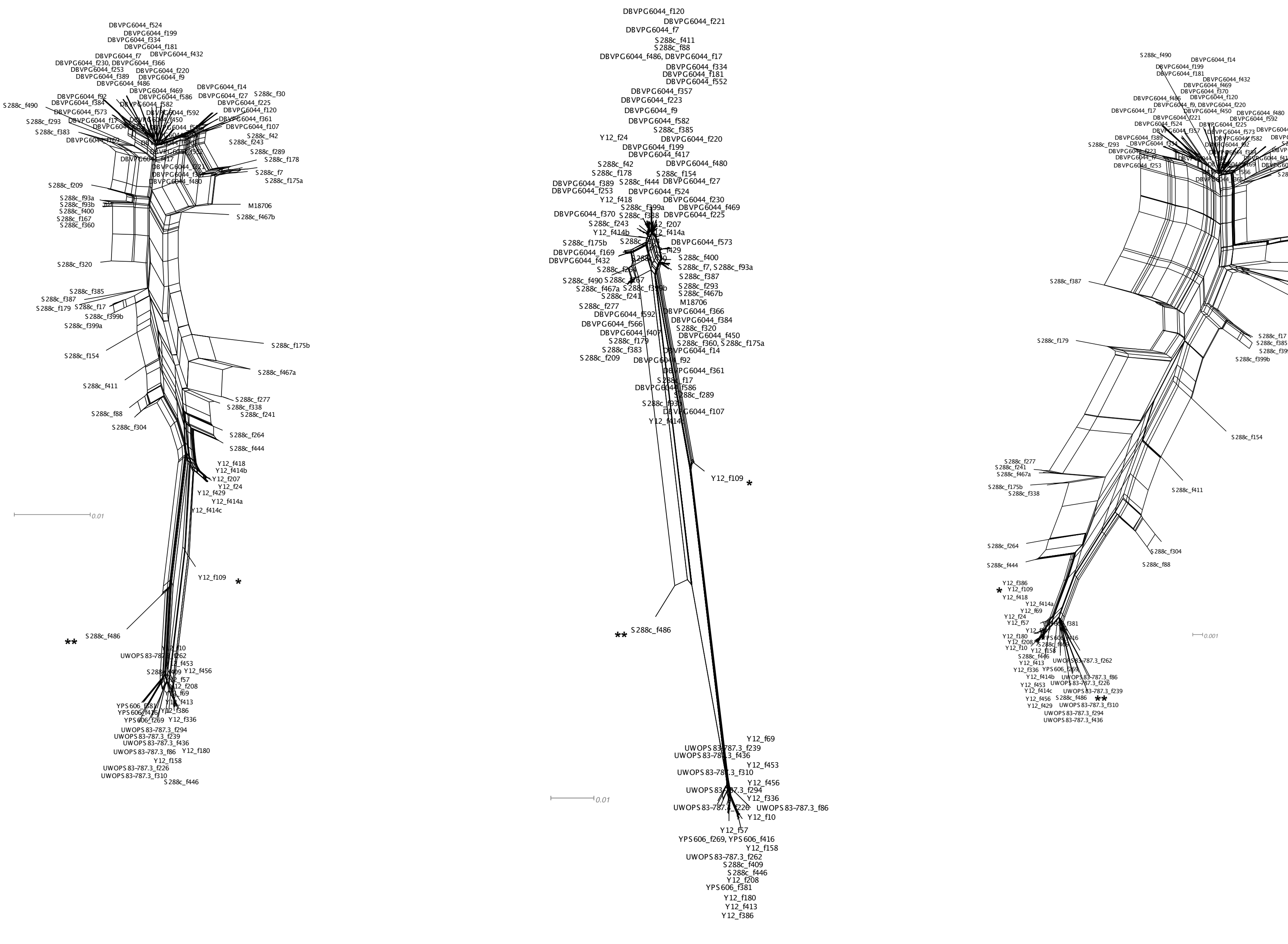

### Supplemental Figure 4

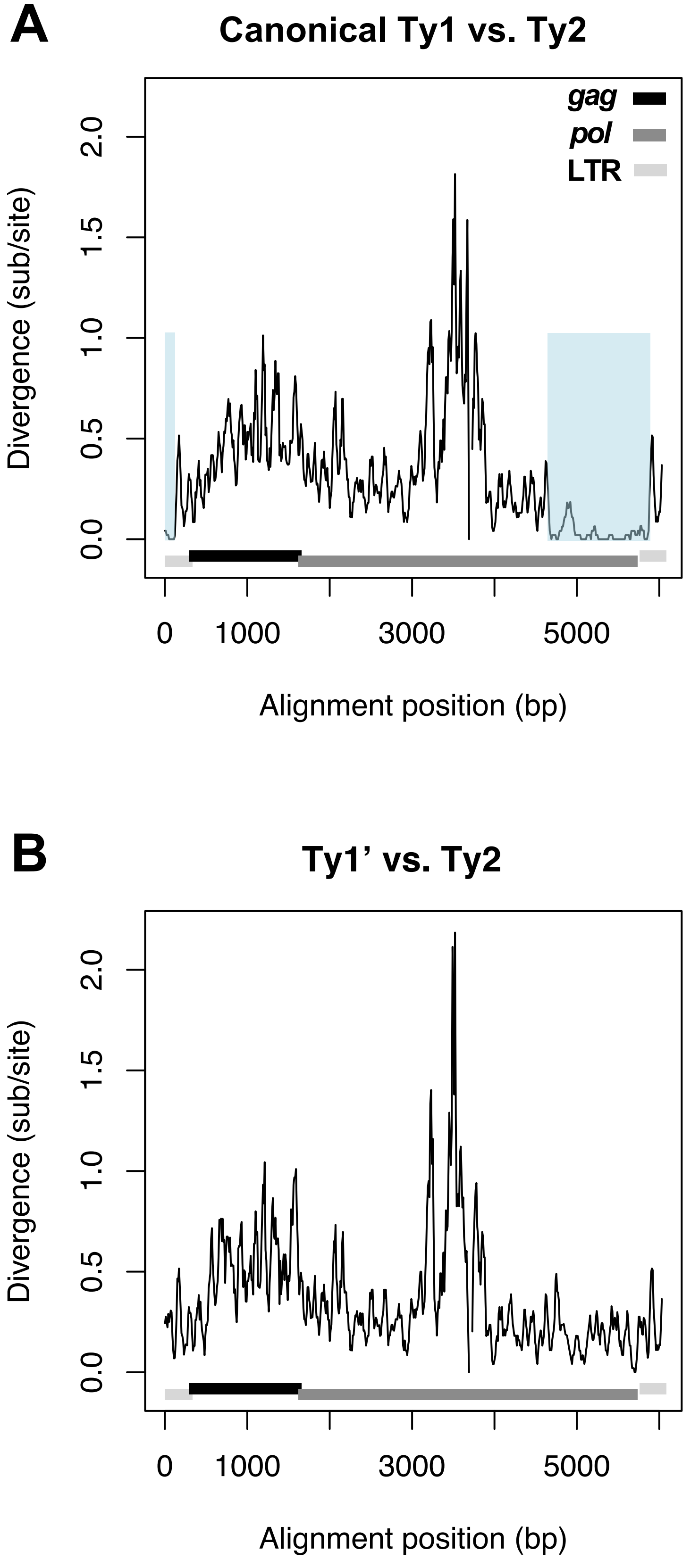

### Supplemental Figure 5

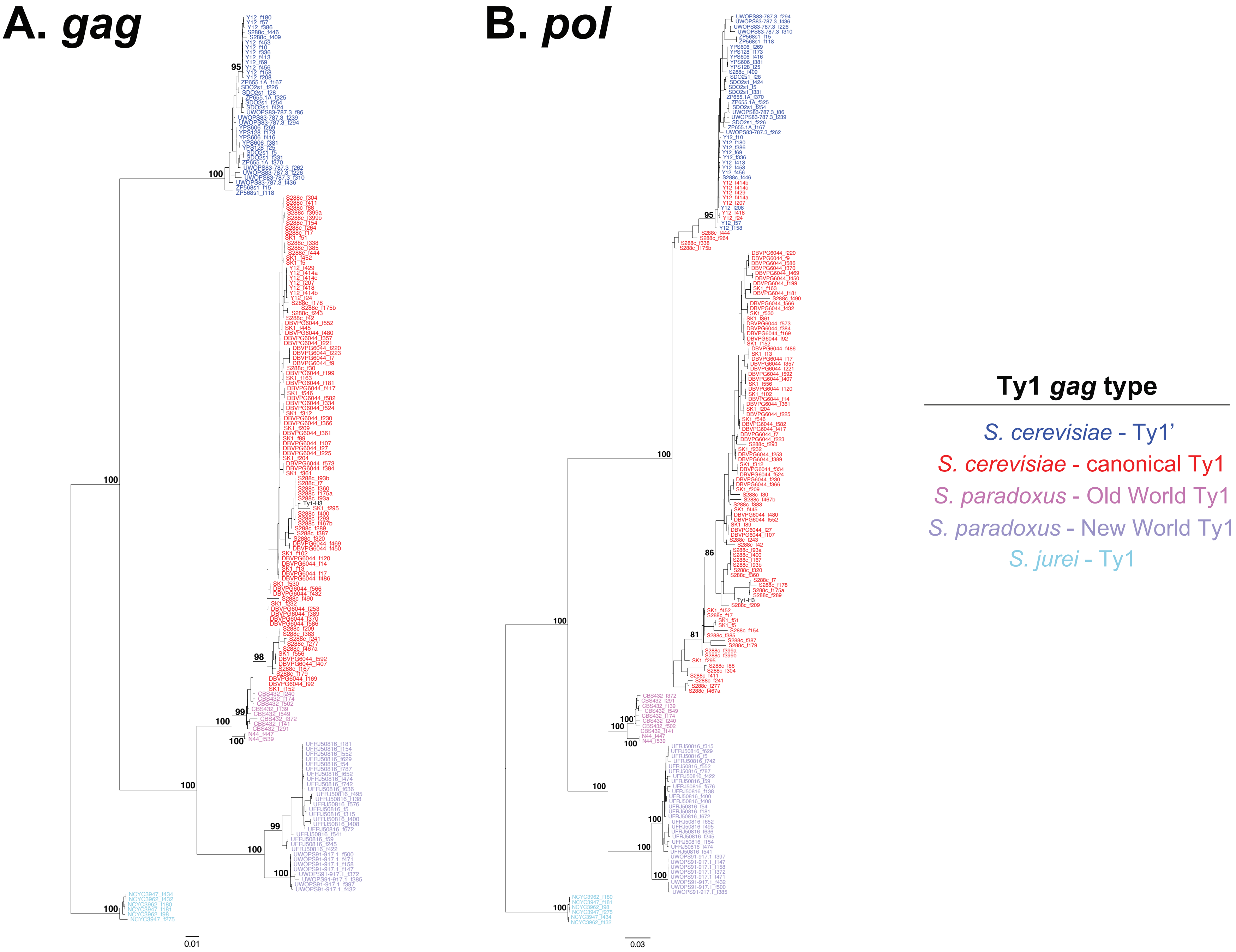

### Supplemental Figure 6

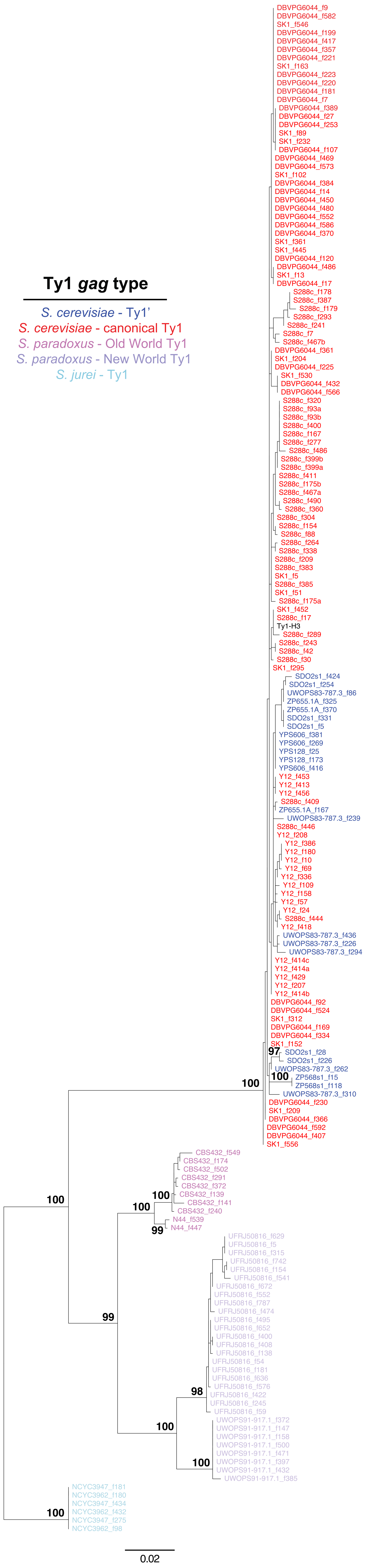
